# The SMARCA5–DMRT1 Pioneer Complex Establishes Epigenetic Priming to Direct Male Germline Development

**DOI:** 10.1101/2025.07.29.667536

**Authors:** Yuka Kitamura, Yasuhisa Munakata, Hironori Abe, Mengwen Hu, Satoyo Oya, Shanmathi Murugesan, Mahnoor Rizwan, Shawna P. Katz, David J. Picketts, Richard M. Schultz, Satoshi H. Namekawa

## Abstract

The establishment of cell type–specific chromatin landscapes is essential for cellular identity, but how these landscapes are generated remains poorly understood. Here, we demonstrate that the chromatin remodeler SMARCA5 establishes epigenetic priming that is required for retinoic acid (RA)–induced differentiation in the male germline. Germ cell–specific deletion of *Smarca5* results in a complete loss of differentiating spermatogonia, phenocopying vitamin A-deficient mice that lack RA signaling. During the perinatal transition from prospermatogonia to undifferentiated spermatogonia, SMARCA5 is recruited to binding sites of the pioneer transcription factor DMRT1, which are located at distal putative enhancers and promoters of germline genes. The SMARCA5–DMRT1 pioneer complex establishes chromatin accessibility at these loci, generating poised enhancers and promoters that serve as RA receptor (RAR)–binding sites. Thus, SMARCA5 licenses transcriptional responses to RA that enable spermatogenic differentiation. Our findings uncover a mechanism linking pioneer factor activity to external signal responsiveness.

**Highlights:** SMARCA5 is required for spermatogonial differentiation

The SMARCA5–DMRT1 pioneer complex establishes epigenetic priming for differentiation

SMARCA5 remodels inaccessible DMRT1-binding sites to an accessible state

SMARCA5 shapes chromatin states that enable retinoic acid responsiveness

## Introduction

The establishment of distinct cellular identities from a shared genetic blueprint requires selective activation of lineage-specific genes and repression of alternative transcriptional programs.^1^ This process is tightly regulated by epigenetic mechanisms, including chromatin remodeling and histone modifications.^2,3^ Emerging evidence suggests that developmental enhancers often acquire accessible chromatin states prior to gene activation, indicating that chromatin preconfiguration can prime regulatory elements for future transcription.^4–6^ However, how such epigenetic priming intersects with differentiation cues—particularly external signals—to direct gene expression programs during development remains poorly understood. Spermatogenesis offers a powerful and experimentally tractable model to investigate this question.

Spermatogenesis is a highly ordered differentiation process driven by a germline stem cell system that supports unidirectional cell differentiation to produce haploid sperm.^7^ This process is maintained by the balanced self-renewal and differentiation of spermatogonial stem cells (SSCs), which reside in a heterogeneous population of slow-cycling, undifferentiated type A spermatogonia (A_undiff_, including A_single_, A_paired_, and A_aligned_ spermatogonia, Figure S1A).^8,9^ Spermatogenic differentiation is orchestrated by retinoic acid (RA) signaling^10–12^ and characterized by dramatic changes in gene expression and epigenetic remodeling.^13,14^ In response to RA signaling, A_undiff_ undergo irreversible commitment to become fast-cycling, KIT^+^ differentiating spermatogonia (designated A1, A2, A3, A4, intermediate (In), and B spermatogonia), which lack stem cell potential.^15^ These spermatogonia undergo a series of mitotic divisions before entering meiosis, ultimately leading to spermiogenesis (Figure 1A). Notably, many genes required for later stages of germ cell differentiation are epigenetically primed in undifferentiated spermatogonia prior to their activation.^16–23^ This finding suggests that epigenetic priming plays a critical role in facilitating RA-responsive transcriptional programs.^24^ Nevertheless, a key unresolved question remains: when and how is epigenetic priming established during the developmental progression that gives rise to undifferentiated spermatogonia?

**Figure 1.**
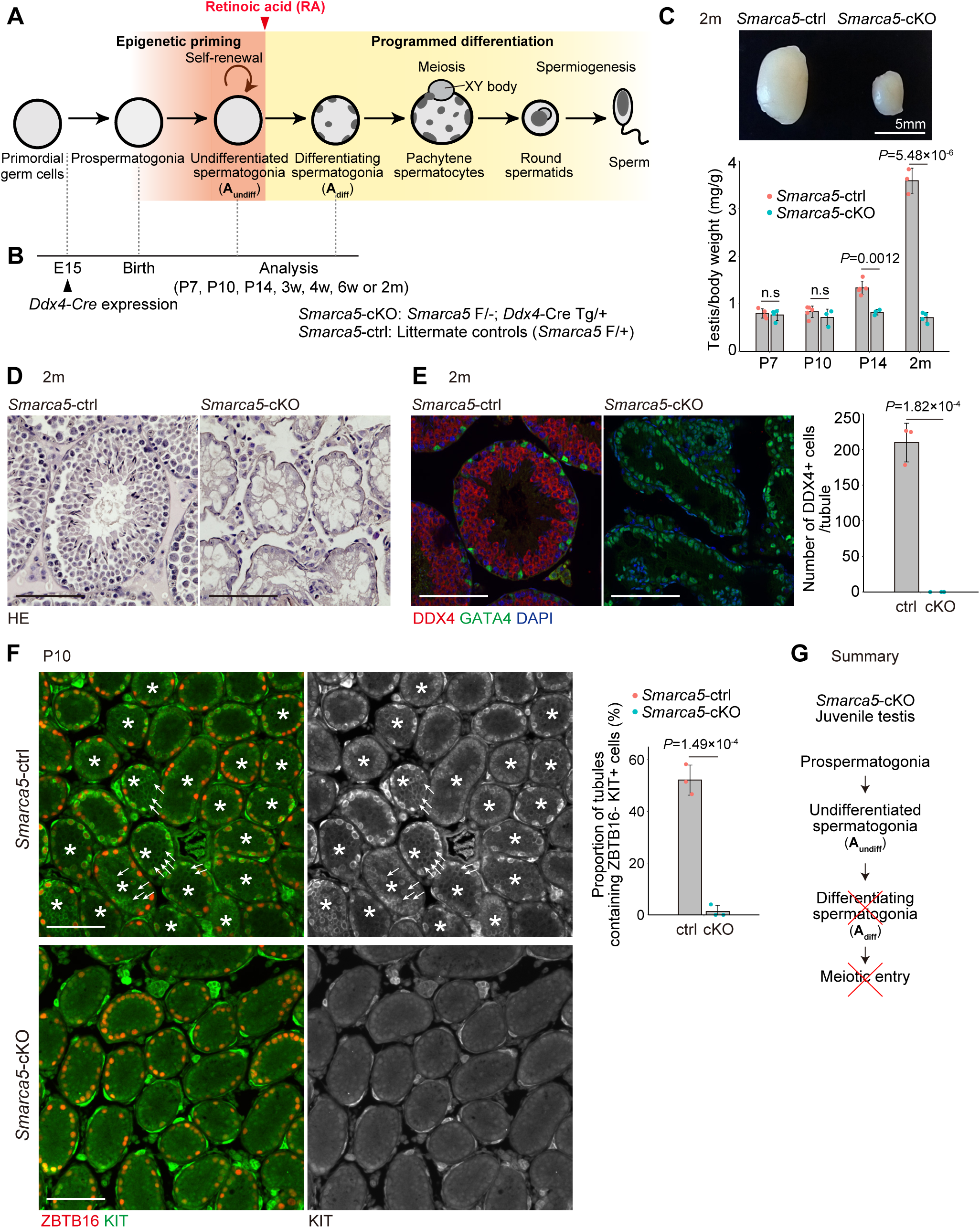
SMARCA5 is required for spermatogonial differentiation. (A) Schematic of the stages of mouse spermatogenesis. (B) Timeline and genotype of the *Smarca5*-cKO used in this study. (C) Image of testes from a *Smarca5*-cKO mouse and a control littermate at 2 months of age. Scale bar, 5 mm. The lower graph shows the testis-to-body weight ratio (mg/g), calculated as the average weight of one testis divided by body weight. Individual data points from each mouse are shown as dots. Error bars represent mean ± standard deviation (s.d.). Statistical significance was assessed using a two-tailed unpaired Student’s t-test with equal variances (n = 3–5 per group). (D) Testis section of *Smarca5*-cKO and control littermate at 2 months, stained with hematoxylin and eosin. Scare bars, 100 μm. (E) Immunofluorescence staining of testis sections from *Smarca5*-cKO and control littermate at 2 months using DAPI and antibodies against DDX4 (germ cell marker) and GATA4 (Sertoli cell marker). Scare bars, 100 μm. The graph shows the number of DDX4⁺ cells per tubule cross-section (19 sections counted per sample). Individual data points represent values from individual mice. Error bars indicate mean ± s.d. Statistical significance was determined by a two-tailed unpaired Student’s t-test assuming equal variances (n = 3 per group). (F) Immunofluorescence staining of testis sections from *Smarca5*-cKO and control littermates at P10 using DAPI and antibodies against ZBTB16 (undifferentiated spermatogonia marker) and KIT (differentiating spermatogonia marker). Asterisks indicate tubule cross-sections containing ZBTB16⁻/KIT⁺ cells, which are marked with arrows. Scale bars, 100 μm. The graph shows the percentage of tubule cross-sections containing ZBTB16⁻/KIT⁺ cells (≥50 cross-sections counted per sample). Individual values represent biological replicates. Error bars indicate mean ± s.d. Statistical significance was determined using a two-tailed unpaired Student’s t-test assuming equal variances (n = 3 per group). (G) Summary of the phenotype observed in juvenile *Smarca5*-cKO mice.

Chromatin accessibility is regulated by two key classes of factors: ATP-dependent chromatin remodelers and pioneer transcription factors. ISWI family remodelers, including SMARCA5 (also known as SNF2H), reposition nucleosomes to generate chromatin environments permissive for transcription factor binding and gene activation.^25–27^ In contrast, pioneer transcription factors can engage their DNA motifs within closed chromatin and initiate local chromatin opening, often enabling recruitment of additional regulatory machinery.^28,29^ Current models posit that pioneer factors recognize nucleosome-occupied target sites and subsequently recruit chromatin remodelers to establish fully-accessible chromatin states.^30^ Although this paradigm is supported by biochemical and *in vitro* cellular reprogramming studies, the *in vivo* relevance and developmental context of this cooperation remain largely unexplored.

Here, we demonstrate that SMARCA5 cooperates with the pioneer transcription factor DMRT1, a critical regulator of male germline development,^31,32^ to establish chromatin accessibility during the perinatal transition from prospermatogonia (also known as gonocytes) to undifferentiated spermatogonia. We show that SMARCA5 is recruited to DMRT1-bound distal regulatory elements and promoters of germline genes and is essential for generating chromatin accessibility at these regions. Despite DMRT1’s intrinsic ability to bind closed chromatin, SMARCA5 is required to remodel these regions into accessible states that enable transcriptional activation of genes critical for spermatogonial maintenance and RA-induced differentiation. These findings uncover a chromatin remodeling mechanism that facilitates the external signal-dependent activation of transcriptional programs essential for spermatogenic differentiation.

## Results

### *Smarca5* knockout in male germ cells causes a complete loss of spermatogonia

To understand how chromatin is primed early in development to support subsequent differentiation, we sought to identify chromatin remodeling mechanisms critical for spermatogenesis. We focused on SMARCA5, which was previously identified in a genetic screen as a key regulator of paternal epigenetic inheritance.^33^ To examine its expression during postnatal spermatogenesis, we reanalyzed single-cell RNA sequencing (scRNA-seq) data^34^ and found that *Smarca5* mRNA is highly expressed from A_undiff_ through to post-meiotic round spermatids (Figure S1B). Consistent with *Smarca5* mRNA expression, SMARCA5 protein was present throughout these stages as well, with particularly high accumulation in A_undiff_ (Figure S1C).

To investigate the role of SMARCA5 in male germ cells, we generated *Smarca5* conditional knockout (*Smarca5*-cKO) mice by crossing mice carrying a *Smarca5* floxed allele^35^ and a germ cell-specific *Ddx4*-Cre transgene, which is expressed from embryonic day 15 (E15)^36^ (Figure 1B). At postnatal day 7 (P7), nearly complete loss of SMARCA5 protein was confirmed in the cKO testis by Western blotting (Figure S1D), and germ cell-specific depletion of SMARCA5 protein was further confirmed by SMARCA5 immunostaining (Figure S1E). Starting at postnatal day 14 (P14), *Smarca5*-cKO mice exhibited reduced testicular size compared to littermate controls (*Smarca5*-ctrl), which carry one deleted and one functional *Smarca5* allele. This reduction became more pronounced and was clearly evident by 2 months of age (Figure 1C). At this stage, the seminiferous tubules in the *Smarca5*-cKO testis were atrophic and devoid of spermatocytes or spermatids (Figure 1D). The absence of germ cells was further confirmed by immunostaining using the germ cell marker DDX4 (Figure 1E). These results indicate that SMARCA5 is essential for spermatogenesis and suggest that the protein is required for both maintenance of spermatogonia and differentiation of male germ cells.

### SMARCA5 is required for spermatogonial differentiation

Because *Smarca5*-cKO testes lack differentiating germ cells, we next investigated whether SMARCA5 is required for the irreversible commitment of A_undiff_ to KIT^+^ differentiating spermatogonia. At postnatal day 10 (P10), when the first-wave spermatogenesis normally reaches the KIT^+^ differentiating spermatogonia stage, *Smarca5*-cKO testes contained cells positive for an A_undiff_ marker ZBTB16 (also known as PLZF), indicating the presence of A_undiff_ (Figure 1F). In contrast, KIT^+^ differentiating spermatogonia were largely absent in *Smarca5*-cKO testis (Figure 1F). Notably, at P10, *Smarca5*-cKO testis largely lacked late differentiating spermatogonia (In and B spermatogonia), characterized by a ZBTB16^-^KIT^+^ profile^37,38^ (Figure 1F).

To further pinpoint the developmental timing of spermatogonia depletion in *Smarca5*-cKO testes, we next examined samples at the earlier stage of P7. During normal spermatogonial development, ZBTB16 is present in early KIT^+^ differentiating spermatogonia (A_1_, A_2_, A_3_, and A_4_, termed differentiating Type A spermatogonia: A_diff_), which have already committed to irreversible differentiation. In *Smarca5*-cKO testes at P7, ZBTB16^+^KIT^+^ A_diff_ were markedly reduced, and late-stage ZBTB16^-^KIT^+^ differentiating spermatogonia were nearly absent, similar to P10. In contrast, the proportion of ZBTB16^+^KIT^-^A_undiff_ was increased compared to littermate *Smarca5*-ctrl testes (Figure S2A). Thus, we conclude that SMARCA5 is required for spermatogonial differentiation (Figure 1G). Supporting this conclusion, we observed that meiosis is not initiated in *Smarca5*-cKO testes, as preleptotene spermatocyte markers STRA8^39^ and MEIOSIN^40^ were absent (Figure S2B).

### SMARCA5 is required for SSC maintenance in the adult testis

The complete absence of germ cells by 2 months of age suggested that maintenance of A_undiff_ is ultimately impaired in *Smarca5*-cKO testis. Indeed, the number of ZBTB16^+^ spermatogonia decreased significantly between 4 and 6 weeks (Figures 2A, B), and during this period, tubules lacking ZBTB16^+^ spermatogonia became apparent (Figure 2C). Therefore, we next sought to determine how SMARCA5 deletion leads to impaired maintenance of A_undiff_. A_undiff_ includes a subpopulation of a relatively small number of A_single_ and A_paired_ cells expressing GFRα1, which form the stem cell pool and give rise to NGN3^+^ cells. However, NGN3^+^ cells can revert to GFRα1^+^ cells, maintaining their long-term self-renewal ability (Figure S1A).^41,42^ In the *Smarca5*-cKO testes, the GFRα1^+^ population was relatively enriched compared to controls at 4 weeks (Figure S2C), likely due in part to the depletion of differentiating spermatogonia. This result further supports the requirement of SMARCA5 for spermatogonial differentiation.

**Figure 2.**
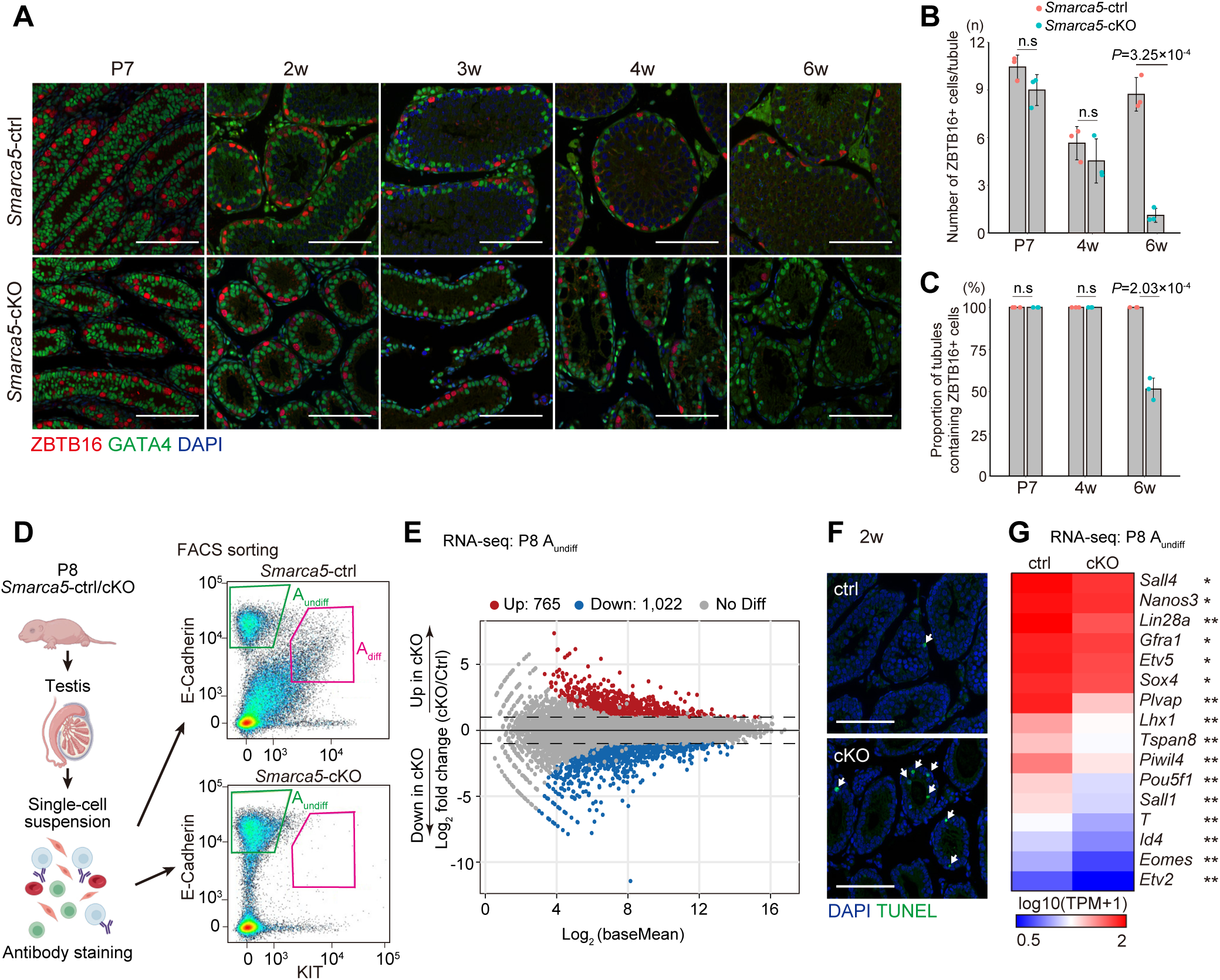
SMARCA5 is required for spermatogonial maintenance in the adult testis. (A) Testis sections of *Smarca5*-cKO and control littermate stained with DAPI and antibodies against ZBTB16 and GATA4 at P7, 2 weeks (2w), 3 weeks (3w), 4 weeks (4w), and 6 weeks (6w). Scare bars, 100 μm. (B) Quantification of ZBTB16⁺ cells per tubule cross-section. At least 30 tubule cross-sections were counted per sample. Error bars represent mean ± standard deviation (s.d.). Statistical significance was determined using a two-tailed unpaired Student’s t-test assuming equal variances (n = 3 per group). (C) Proportion of tubule cross-sections containing ZBTB16⁺ cells. Error bars represent mean ± s.d. Statistical significance was determined using a two-tailed unpaired Student’s t-test assuming equal variances (n = 3 per group). (D) Left: Schematic overview of the experimental workflow for isolating A_undiff_ and A_diff_. Right: FACS profiles showing sorting of A_undiff_ (green gate) and A_diff_ (pink gate) populations from *Smarca5*-control and -cKO testes. (E) Transcriptome comparison between *Smarca5*-ctrl and cKO A_undiff_. Differentially expressed genes (DEGs) are defined as those with Log_2_ fold change > 2, *Padj* < 0.05, based on a binomial test with Benjamini-Hochberg correction. (F) TUNEL staining of testis sections from *Smarca5*-cKO and control littermates at 2 weeks. TUNEL⁺ cells are indicated by arrows. Scare bars, 100 μm. (G) Heatmap showing gene expression of representative A_undiff_ markers in *Smarca5-*ctrl and cKO A_undiff_. *: *Padj* < 0.05, **: *Padj* < 0.05 and Log_2_ fold change > 1.

We next examined whether *Smarca5*-cKO A_undiff_ exhibited an abnormal cell cycle profile, as slow cycling is essential for maintenance of A_undiff_. To this end, we assessed the proportion of active cycling A_undiff_ cells at 2 and 4 weeks by immunostaining with Ki67, a proliferation marker present in G1, S, G2, and M phases of the cell cycle, but absent in G0.^43^ Consistent with the slow cycling nature of A_undiff_, only 22.5% of A_undiff_ in *Smarca5-*ctrl mice were Ki67^+^ at 2 weeks (Figure S2D). However, 83.3% of A_undiff_ in *Smarca5*-cKO testes were Ki67^+^, an ∼ 3-fold increase compared to control testes (Figure S2D). This suggests that *Smarca5*-cKO A_undiff_ are highly proliferative at 2 weeks. At 4 weeks, the proportion of Ki67^+^cells remained high in *Smarca5*-cKO A_undiff_. These findings indicate that SMARCA5 is required to maintain the slow-cycling state of A_undiff_, and that their overproliferation in *Smarca5*-cKO testes ultimately leads to depletion of the SSC pool.

### SMARCA5 promotes expression of SSC maintenance genes

To further determine the cause of SSC maintenance defects, we examined gene expression profiles by performing RNA-sequencing (RNA-seq) on spermatogonia at P8, a stage when sufficient numbers of both A_undiff_ and A_diff_ can be obtained from normal testes. We isolated A_undiff_ and A_diff_ from *Smarca5*-ctrl males at P8 using our previously established fluorescence-activated cell sorting (FACS) method (Figure 2D).^8,44^ We utilized the cell surface marker E-Cadherin, which is expressed in both A_undiff_ and A_diff_, to isolate these populations, and employed KIT expression to specifically identify A_diff_ (Figure 2D). In addition, we used the cell surface marker CD9 to exclude somatic contaminants^45^, thus ensuring the isolation of a highly pure spermatogonial population. Using this method, A_undiff_ were collected based on E-cadherin and high CD9 expression, and absence of KIT (E-cadherin^+^CD9^high^KIT^−^), while A_diff_ were isolated based on E-cadherin and KIT expression with medium CD9 level (E-cadherin^+^CD9^medium^KIT^+^). As expected based on our immunostaining results from the P7 testes (Figure S2A), at P8, *Smarca5*-cKO males lacked an A_diff_ population (Figure 2D). We confirmed the high purity of A_undiff_ and A_diff_ (Figure S3A) and observed strong similarity between biological replicates in the RNA-seq data (Figure S3B).

During normal spermatogonial differentiation from A_undiff_ to A_diff_ at P8, a major shift in gene expression occurred, with 2,639 genes upregulated and 1,559 genes downregulated (Figure S3C). However, in *Smarca5-*cKO A_undiff_, the overall gene expression profile was distinct from both *Smarca5*-ctrl A_undiff_ and A_diff_ (Figure S3B). Compared to *Smarca5-*ctrl A_undiff_, *Smarca5-*cKO A_undiff_ showed 765 upregulated genes and 1,022 downregulated genes (Figure 2E). Gene ontology (GO) analysis revealed that the upregulated genes in *Smarca5-*cKO A_undiff_ were enriched for apoptosis-related terms (Figure S3D). Consistent with this, the frequency of apoptotic cell death was increased in *Smarca5-*cKO testes (Figure 2F).

Loss of SMARCA5 in A_undiff_ led to the ectopic upregulation of genes that were normally upregulated from A_undiff_ to A_diff_ (termed “A_diff_-high genes”: Figure S3E, F). Conversely, genes that were highly expressed in A_undiff_, which are typically downregulated in A_diff_ (termed “A_undiff_-high genes”), were downregulated in *Smarca5*-cKO A_undiff_ (Figure S3E, G). These downregulated genes in *Smarca5-*cKO, enriched for the GO term “regulation of transcription by RNA polymerase II” (Fig. S3D), which was also the top-ranked GO term among A_undiff_ high genes (Fig. S3E). These findings indicate that *Smarca5*-cKO A_undiff_ exhibit impaired expression of A_undiff_-specific genes. Consistent with this, key SSC maintenance genes —such as *Sall4*, *Nanos3*, *Lin28a, Gfra1, Etv5, Sox4, and Plvap*—were down-regulated upon *Smarca5* loss (Figure 2G), indicating that SMARCA5 is required for proper expression of SSC maintenance genes. Together, these results suggest that *Smarca5*-cKO A_undiff_ display upregulation of apoptosis-related genes and downregulation of SSC maintenance genes, which may ultimately lead to A_undiff_ depletion in adult testes.

### SMARCA5 deficiency results in a closed chromatin state at DMRT1-binding sites

As SMARCA5 is an ATP-dependent chromatin remodeler,^46^ we next investigated whether SMARCA5 establishes the chromatin landscape required for SSC maintenance and spermatogonial differentiation. To this end, we performed an assay for transposase-accessible chromatin using sequencing (ATAC-seq)^47,48^ to assess genome-wide chromatin accessibility in A_undiff_ and A_diff_ from *Smarca5-*ctrl mice and A_undiff_ from *Smarca5*-cKO mice at P8. Due to the absence of A_diff_ in *Smarca5*-cKO testes, we were unable to analyze this population. Pearson correlation coefficient analysis confirmed high correlations among biological replicates (Figure S4A). Among 17,647 ATAC peaks detected in *Smarca5-*ctrl A_diff_, 11,290 peaks (64.0%) overlapped with those in A_undiff_ (Figures S4B, C), and we identified 6,203 A_undiff_-specific peaks and 6,357 A_diff_-specific peaks. This suggests that a large core set of accessible chromatin regions is shared between the two stages, although chromatin accessibility at some specific regions shifts during the transition from A_undiff_ to A_diff_ (Figure S4C). Many of the ATAC peaks in A_undiff_ were located within 10 kb of genes expressed in A_undiff_ (e.g., *Etv2*, *Etv5*, *Id4*, and *Ret*) and A_diff_ (e.g., *Dmc1*, *Meioc*, *Stra8*, and *Prdm9*) (Figure S4D), suggesting that A_undiff_ ATAC peaks serve as proximal or distal regulatory elements for both A_undiff_ and A_diff_. Taken together, A_undiff_ and A_diff_ exhibit largely overlapping chromatin accessibility profiles, with putative regulatory elements of genes upregulated in A_diff_ already accessible at the A_undiff_ stage. These findings suggest that the gene expression program for spermatogonial differentiation is primed in A_undiff_.

Notably, A_undiff_ from *Smarca5*-cKO mice exhibited a largely distinct chromatin accessibility compared to A_undiff_ and A_diff_ from *Smarca5-*ctrl mice (Figure S4A). In A_undiff_, loss of SMARCA5 significantly affected the number of ATAC peaks, especially at the intergenic and intronic regions (Figure 3A). In contrast, ATAC peaks at promoter regions tended to be common between *Smarca5*-ctrl and cKO. These results suggest that SMARCA5 primarily regulates chromatin accessibility at distal regulatory regions, such as enhancers, rather than at promoters.

**Figure 3.**
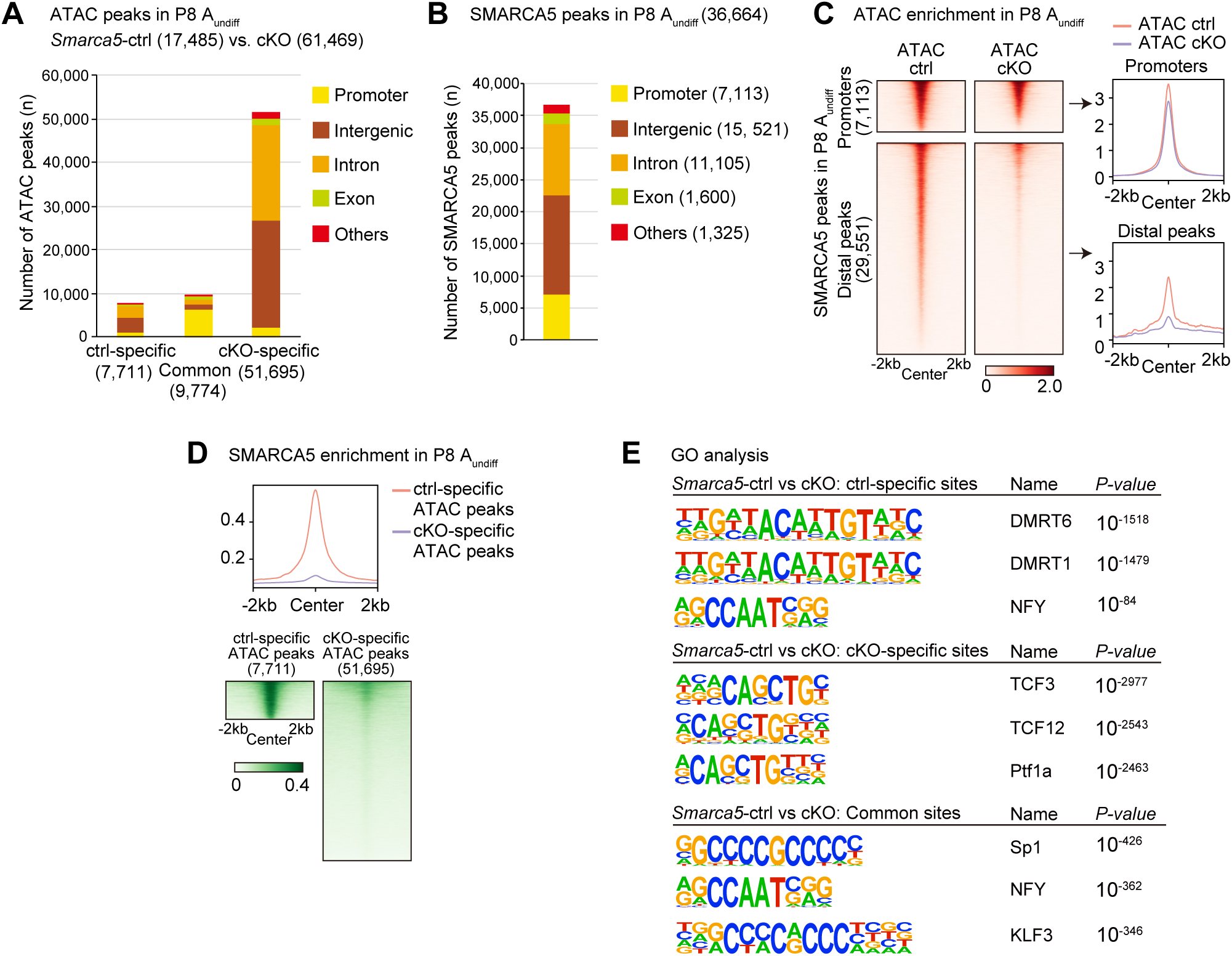
SMARCA5 directly promotes chromatin accessibility at its binding sites, predominantly at distal regulatory regions. (A) Numbers and genomic distribution of ATAC-seq peaks in *Smarca5*-ctrl and cKO A_undiff_. (B) Numbers and genomic distribution of SMARCA5 CUT&Tag peaks in A_undiff_. (C) Heatmaps and average tag density plots showing ATAC enrichment in *Smarca5*-ctrl and cKO A_undiff_ at promoter regions (top) and distal SMARCA5-binding peaks (bottom). (D) Heatmaps and average tag density plots showing SMARCA5 enrichment at *Smarca5*-ctrl-specific ATAC peaks or *Smarca5*-cKO-specific ATAC peaks. (E) HOMER known motif analyses of ATAC-seq peaks at *Smarca5*-ctrl-specific, cKO-specific, and common sites.

To determine if SMARCA5 directly binds to chromatin regions with altered accessibility in *Smarca5*-cKO, we performed Cleavage Under Targets and Tagmentation (CUT&Tag)^49^ analysis of SMARCA5. The majority of SMARCA5 peaks were located in intergenic (42.3%) and intronic (30.3%) regions, and 19.4% were found at promoter regions (Figure 3B). These SMARCA5 binding sites were largely accessible at both promoter regions (within ± 1 kb from transcription start sites, TSSs) and distal regions (beyond ± 1 kb from TSSs) in *Smarca5*-ctrl cells. Whereas promoter accessibility was retained in *Smarca5*-cKO, distal SMARCA5 peaks lost accessibility (Figure 3D). SMARCA5 was also enriched at ATAC peaks specific to *Smarca5*-ctrl, but not at ATAC peaks specific to *Smarca5*-cKO (Figure 3E). These findings indicate that SMARCA5 directly promotes chromatin accessibility at its binding sites, predominantly at distal regulatory regions. While *Smarca5*-cKO also leads to the emergence of ectopically accessible regions, the near absence of SMARCA5 binding at these sites suggests that many of them are indirectly affected by the loss of SMARCA5.

We further investigated the characteristics of ATAC peaks specific to *Smarca5*-ctrl, as these are presumably direct targets of SMARCA5. Motif analysis of ATAC peaks using HOMER revealed that motifs shared by the transcription factors DMRT1 and DMRT6 were enriched in *Smarca5*-ctrl A_undiff_-specific peaks (Figure 3E). Interestingly, DMRT1 is expressed in male germ cells postnatally and present in A_undiff_, ^50^ regulates SSC maintenance^51^ and suppresses precocious meiotic entry in spermatogonia^52^. Thus, SMARCA5 might collaborate with DMRT1 to promote the establishment of chromatin states conducive to male germ cell development. We focused on DMRT1 rather than DMRT6 due to their distinct expression profiles; DMRT1 is expressed in male germ cells postnatally and is present in A_undiff_, ^50^ whereas DMRT6 expression begins at the A_diff_ stage and functions during spermatogenic differentiation.^53^ Therefore, DMRT6 is unlikely to be involved in SMARCA5-regulated distal accessible regions in A_undiff_. To further investigate the possible role of DMRT1, we performed a CUT&Tag analysis for DMRT1 in A_undiff_ (Figure S4E). DMRT1 peaks were predominantly located at intergenic (42.4%) and intronic (35.8%) regions, while 17.8% of the DMRT1 peaks were located in promoter regions (Figure 4A). Compared to previous DMRT1 ChIP-seq studies using whole testis,^51^ our CUT&Tag analysis identified a greater number of promoter-associated peaks. Nonetheless, consistent with the previous study^51^, DMRT1 exhibited stronger binding at distal regions compared to promoters (Figure 4B). While DMRT1-bound promoter regions remained accessible in *Smarca5*-cKO, distal DMRT1 peak regions showed significantly reduced accessibility upon SMARCA5 loss (Figure 4B). These findings indicate that SMARCA5 promotes chromatin accessibility at distal DMRT1-binding sites.

**Figure 4.**
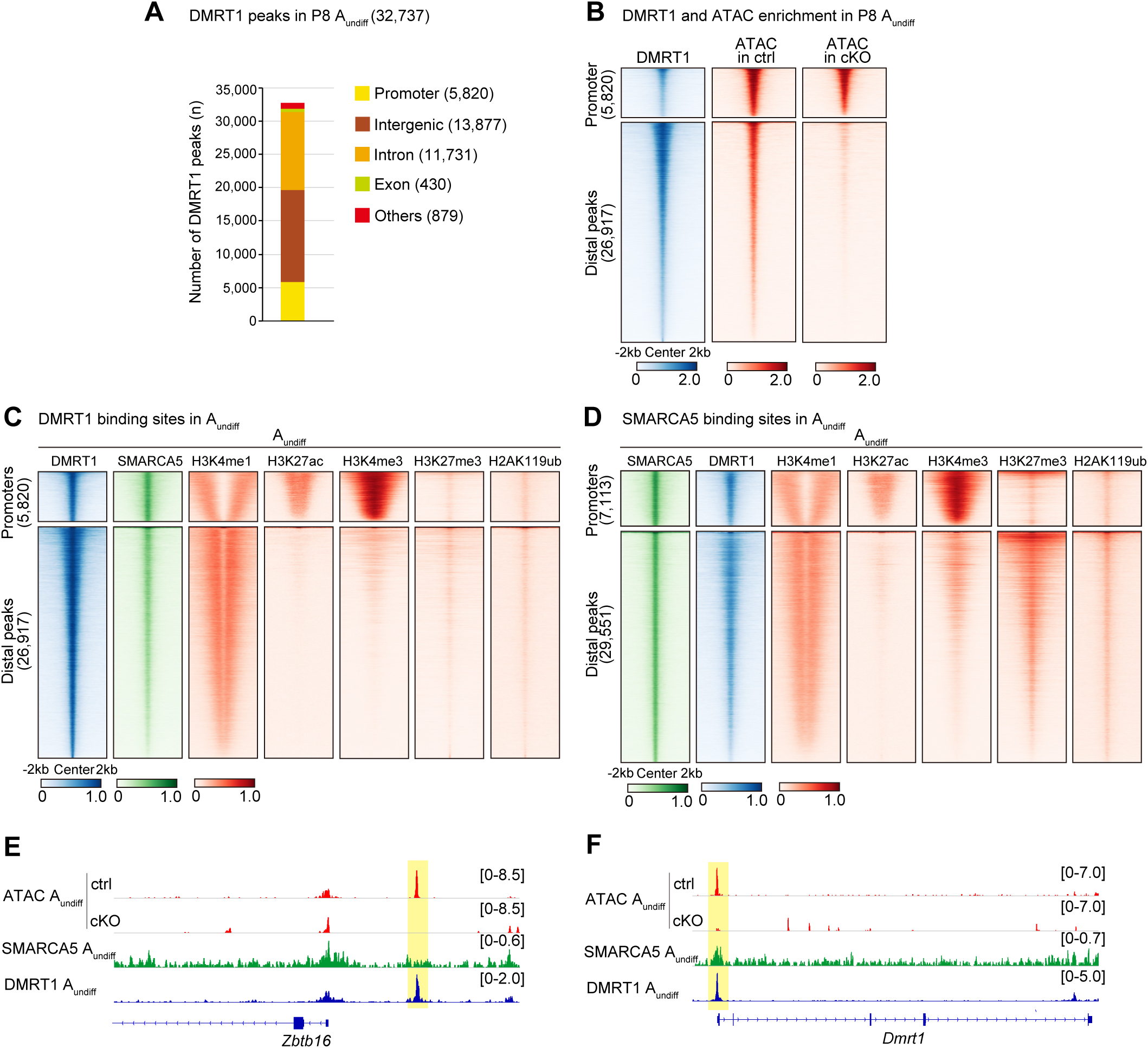
SMARCA5 establishes chromatin accessibility at DMRT1-binding sites. (A) Number and genomic distribution of DMRT1 CUT&Tag peaks in A_undiff_. (B) Heatmaps showing DMRT1 CUT&Tag and ATAC enrichment in *Smarca5*-ctrl A_undiff_ and *Smarca5*-cKO A_undiff_ at promoter regions and distal DMRT1-binding peaks. (C and D) Heatmaps showing enrichment of the indicated factors (DMRT1, SMARCA5, H3K4me1, H3K27ac, H3K4me3, H3K27me3, and H2AK119ub) in A_undiff_ at promoter regions and distal DMRT1-binding peaks (C) or distal SMARCA5-binding peaks (D). (E and F) Track view of *Zbtb16* (E) and *Dmrt1* (F) gene loci showing ATAC peaks in *Smarca5*-ctrl and cKO A_undiff_ and CUT&Tag profiles for SMARCA5 and DMRT1 in *Smarca5*-ctrl A_undiff_. Regions with reduced ATAC signal upon *Smarca5* deletion—corresponding to a putative enhancer of *Zbtb16* (E) and the *Dmrt1* promoter (F)—are highlighted in yellow.

### SMARCA5 establishes chromatin accessibility at DMRT1-binding sites

We next sought to elucidate the functions of distal regulatory elements regulated by SMARCA5 and DMRT1. To this end, we reanalyzed publicly available chromatin profiling datasets from A_undiff_ to examine the distribution of representative histone modifications at these distal regions.^44,54^ We found that H3K4me1 (monomethylation of histone H3 at lysine 4), a hallmark of poised enhancers, was enriched at distal DMRT1 peaks. These regions lacked the active enhancer/promoter mark H3K27ac (H3K27 acetylation)^20^ and the promoter mark H3K4me3^55^ (Figure 4C), suggesting that distal elements are poised rather than active enhancers. Additionally, these sites retained repressive histone modifications, including H2AK119ub and H3K27me3^44^, which are mediated by Polycomb repressive complexes PRC1 and PRC2, respectively. This indicates a role for Polycomb in enhancer poising. Notably, H3K4me1 was enriched specifically at regions adjacent to the peak centers. In contrast, DMRT1-associated promoters showed strong enrichment of both H3K27ac and H3K4me3, consistent with an active promoter status (Figure 4C). Analysis of SMARCA5 binding sites in A_undiff_ revealed similar enrichment of H3K4me1 (Figure 4D). A subset of these SMARCA5-bound distal regions also displayed H2AK119ub and H3K27me3 marks. Overall, these results suggest that SMARCA5 and DMRT1 largely co-occupy putative poised enhancers, with SMARCA5-bound regions exhibiting a stronger association with Polycomb-mediated repression than those bound by DMRT1.

We further investigated how individual loci are regulated by SMARCA5 and DMRT1. DMRT1 is known to bind a putative upstream enhancer of the *Zbtb16* locus and regulate its expression^51^. We found that chromatin accessibility at this enhancer was also SMARCA5-dependent and that both SMARCA5 and DMRT1 bound to this region (Figure 4E), suggesting that they act cooperatively to activate *Zbtb16* expression. In adult A_undiff_ from *Smarca5*-cKO mice, we observed reduced expression of the ZBTB16 protein (Figure S4G), a phenotype resembling that of *Dmrt1*-deficient spermatogonia^51^. Additionally, SMARCA5 and DMRT1 co-bound the *Dmrt1* promoter, where chromatin accessibility was likewise SMARCA5-dependent (Figure 4F), suggesting that SMARCA5 and DMRT1 cooperate to establish the chromatin landscape at the *Dmrt1* promoter. Together, these findings support a model in which SMARCA5 and DMRT1 work in concert to establish chromatin accessibility at key regulatory elements critical for spermatogenesis.

### SMARCA5 establishes chromatin accessibility at DMRT1-binding sites after the prospermatogonia stage

A key outstanding question is when SMARCA5 establishes chromatin accessibility at DMRT1-binding sites in the male germline. To address this, we performed ATAC-seq on prospermatogonia (ProSG) at P0 (Figure S5A) and examined changes in chromatin accessibility during the transition to spermatogonia. In *Smarca5*-control P0 ProSG, we identified 12,780 ATAC peaks, approximately one-third of which were located at promoter regions (Figure 5A). From P0 ProSG to P8 A_undiff_, there was a progressive increase in ATAC peaks (Figure 5B). The P8 A_undiff_-specific peaks that emerged during this developmental window were enriched in the DMRT1/6 motif (Figures S5B). When comparing the accessible chromatin landscapes of P0 ProSG and P8 A_undiff_, we found that stage-specific ATAC peaks were predominantly located in intergenic and intronic regions, whereas more than half of the shared peaks were found at promoter regions (Figure 5C). The newly acquired ATAC peaks in A_undiff_ were enriched for the poised enhancer mark H3K4me1 (Figure 5D). Based on these findings, we hypothesized that SMARCA5 actively generates chromatin accessibility at these poised enhancers. Indeed, distal DMRT1-binding sites in P8 A_undiff_ gained chromatin accessibility during this transition, and this gain was dependent on SMARCA5 (Figure 5G). These findings indicate that SMARCA5 establishes chromatin accessibility at distal elements of DMRT1-binding sites during the transition from P0 ProSG to P8 A_undiff_.

**Figure 5.**
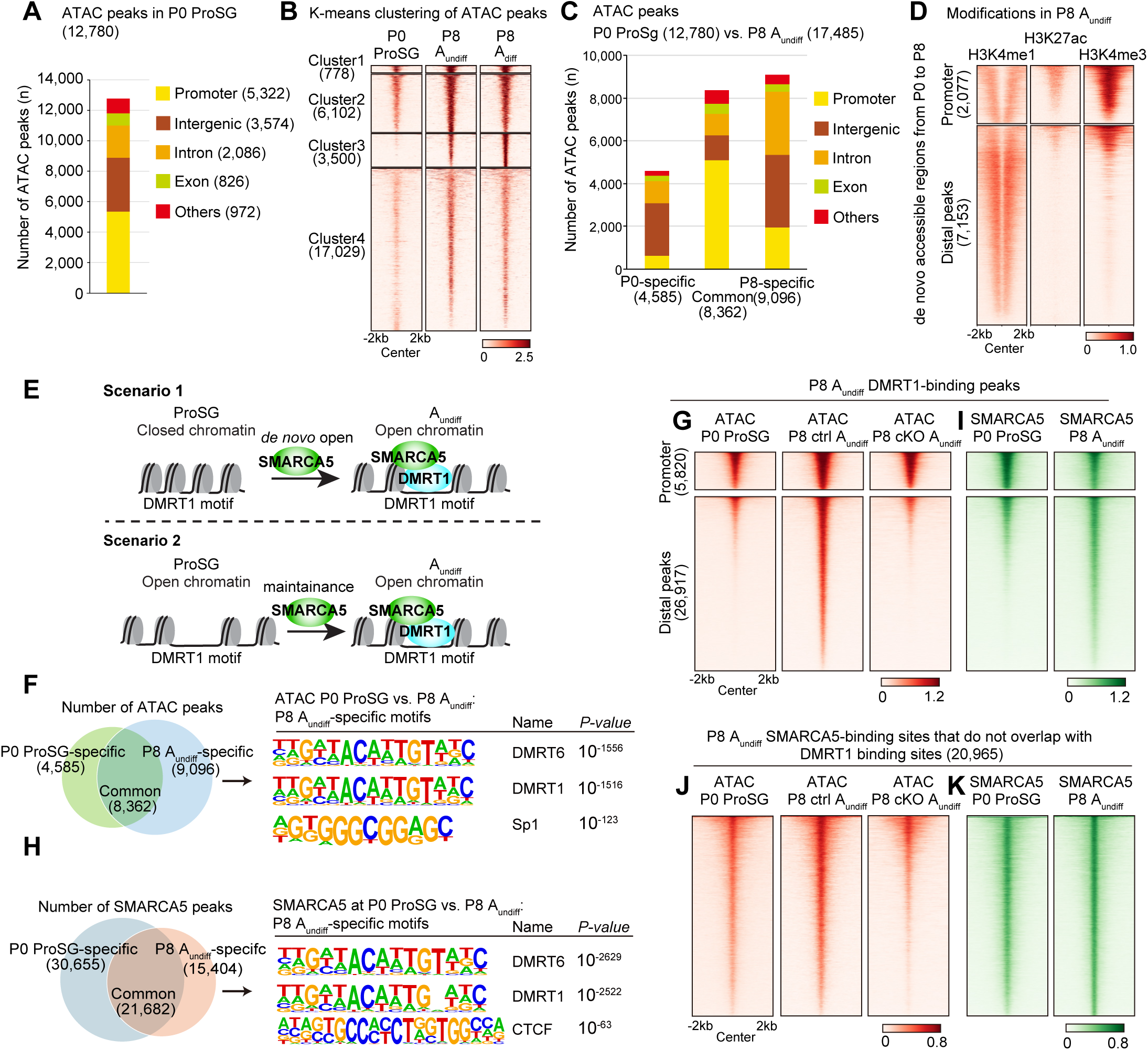
SMARCA5 is recruited to distal regions of DMRT1-binding sites during the transition from ProSG to A_undiff_. (A) Number and genomic distribution of ATAC peaks in P0 ProSG. (B) Heatmap of k-means clustering showing progressive gain of chromatin accessibility from ProSG to A_undiff_ and A_diff_. (C) Number and genomic distribution of ATAC-seq peaks in P0 ProSG and P8 A_undiff_. (D) Heatmaps showing enrichment of H3K4me1, H3K27ac, and H3K4me3 in A_undiff_ at promoter regions and distal peaks of de novo accessible region acquired during the ProSG-to-A_undiff_ transition. (E) Schematic model illustrating potential mechanisms underlying chromatin state changes during the ProSG-to-A_undiff_ transition. (F and H) Left: Venn diagram comparing ATAC-seq peaks (F) and SMARCA5 peaks (H) in ProSG and A_undiff_. Right: HOMER known motif analysis of A_undiff_-specific ATAC-seq peaks (F) and SMARCA5 peaks (H). (G and I) Heatmaps showing ATAC enrichment (G) and SMARCA5 enrichment (I) at promoter regions and distal DMRT1-binding peaks identified in A_undiff_.(J and K) Heatmaps showing ATAC enrichment (J) and SMARCA5 enrichment (K) at SMARCA5-binding sites that do not overlap with DMRT1-binding sites in A_undiff_.

### SMARCA5 is recruited to distal DMRT1-binding sites to establish accessible chromatin

To determine when SMARCA5 is recruited to distal DMRT1-binding sites, we performed CUT&Tag for SMARCA5 in P0 ProSG (Figure S5C) and compared the binding profiles to P8 A_undiff_. We observed that 30,655 P0 ProSG SMARCA5 peaks were lost, while 15,404 new SMARCA5 peaks were gained in P8 A_undiff_ (Figure 5H). Notably, DMRT1/6 motifs were enriched within the P8-specific SMARCA5 peaks (Figure 5H, S5D). At DMRT1-binding sites in P8 A_undiff_, SMARCA5 was newly recruited to many distal regions, whereas it was already bound to promoter regions (Figure 5I). This pattern was specific to DMRT1-binding sites, as SMARCA5 peaks in A_undiff_ that did not overlap with DMRT1-binding sites were already accessible and SMARCA5-bound in ProSG (Figures 5J, K). Consistent with these findings, SMARCA5 was expressed in germ cells from E18.5 through P3 (Figure S5E). In contrast, DMRT1 was not expressed at E18.5 but appeared in a subset of germ cells at P0 and was present in most germ cells by P3 (Figure S5F). Thus, the onset of DMRT1 expression (from P0 onwards) coincided with the *de novo* establishment of accessibility at distal DMRT1-binding sites, suggesting that SMARCA5 recognizes distal DMRT1-binding sites upon DMRT1 expression and facilitates the generation of accessible chromatin at these sites.

### The SMARCA5–DMRT1 pioneer complex binds closed chromatin and facilitates DNA accessibility

A previous study suggested that DMRT1 functions as a pioneer transcription factor, capable of binding closed chromatin to facilitate chromatin accessibility.^56^ Based on our findings, we hypothesized that DMRT1 cooperates with SMARCA5 to generate accessible chromatin at distal DMRT1-binding sites. To test this, we assessed DMRT1’s chromatin-binding ability using CUT&Tag in P8 A_undiff_ from *Smarca5*-cKO mice (Figure S5G) and compared it to control mice (Figure S4F). DMRT1 enrichment at its binding sites was largely similar between *Smarca5*-ctrl and cKO, with minimal differences observed at both promoter and distal regions (Figure 6A). Notably, DMRT1 binding was retained at sites that remained closed in *Smarca5*-cKO but were accessible in controls—specifically at SMARCA5-dependent ATAC peaks (Figure 6B). These results indicate that DMRT1 is capable of binding closed chromatin and that SMARCA5 is required to remodel these regions into an accessible state at distal DMRT1-binding sites.

**Figure 6.**
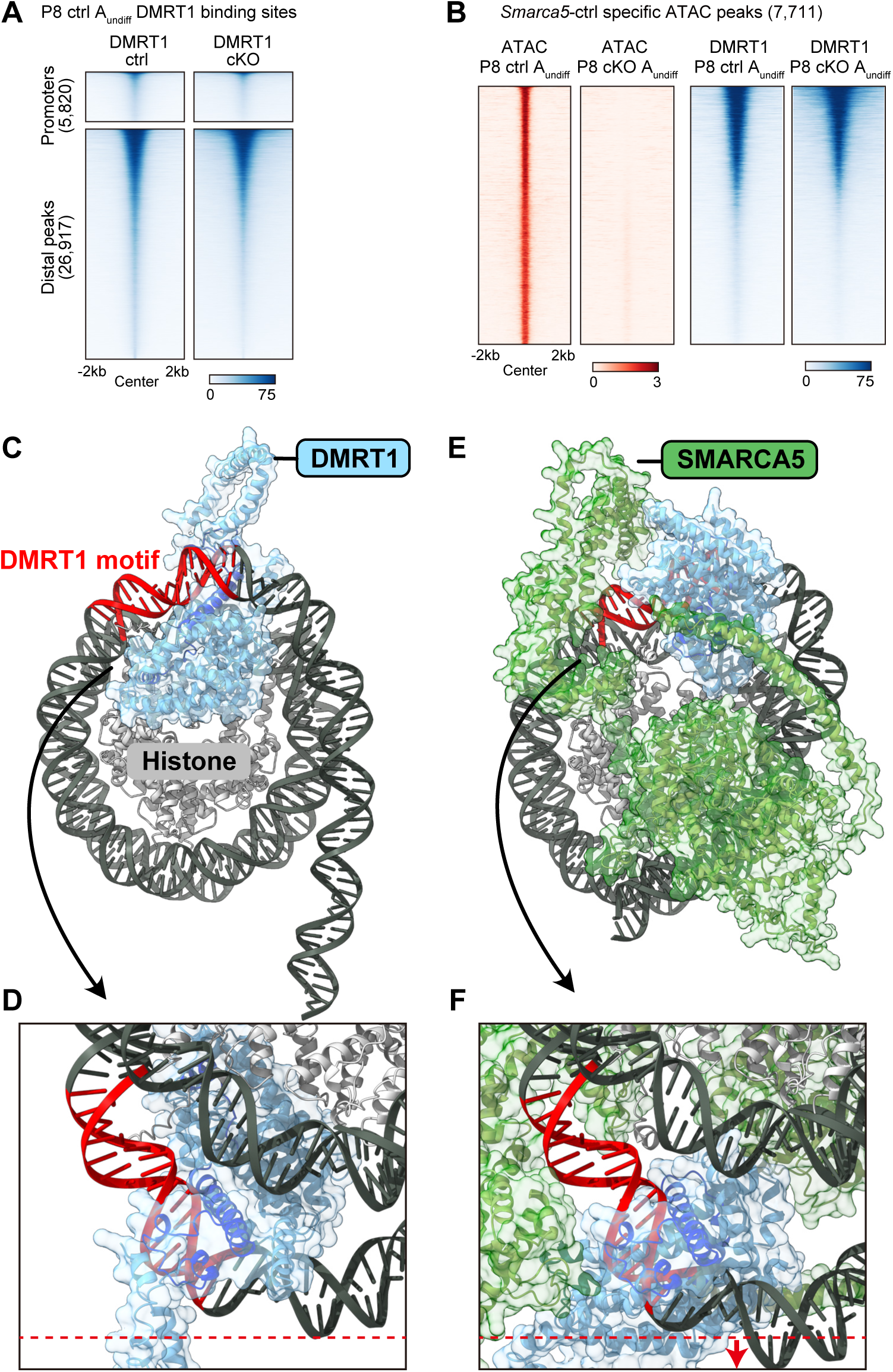
The SMARCA5-DMRT1 pioneering complex establishes chromatin accessibility. (A) Heatmaps showing DMRT1 CUT&Tag signal in *Smarca5*-ctrl and *Smarca5*-cKO A_undiff_-at promoter regions and distal DMRT1-binding peaks identified in A_undiff_. (B) Heatmaps showing ATAC-seq and DMRT1 enrichment in *Smarca5*-ctrl and *Smarca5*-cKO A_undiff_ at ATAC peaks specific to *Smarca5*-ctrl A_undiff_. Signal intensities (bar plots below the heatmaps) are normalized using BPM (Bins Per Million mapped reads) for ATAC-seq and RPKM (Reads Per Kilobase per Million mapped reads) for DMRT1 CUT&Tag. (C–F) Structural predictions of DMRT1–nucleosome (C, D) and DMRT1–SMARCA5– nucleosome (E, F) interactions using AF3. To compare the distance between the histone core and DNA, the region of DNA furthest from the histone core is marked with a red dotted line in (D). In (F), red arrows indicate the displacement of DNA in the presence of SMARCA5, suggesting chromatin loosening upon complex formation.

We modeled chromatin relaxation by DMRT1 with and without SMARCA5 using AlphaFold3 (AF3), which enables highly accurate predictions of protein–nucleic acid complexes.^57^ We used the 601 DNA sequence, commonly employed for nucleosome modeling^58^ and incorporated a DMRT1-binding motif. Structural predictions were then generated for DMRT1 alone and for the DMRT1–SMARCA5 complex. The DM domain of DMRT1, known to interact with DNA,^59^ was predicted to bind its motif via a major α-helix (Figure 6C, D). This binding mode closely matches the structure of the human DMRT1– DNA complex determined by X-ray crystallography.^60^ Notably, AF3 predicts that DMRT1 can bind DNA wrapped around histones, consistent with our CUT&Tag data showing DMRT1 occupancy at inaccessible chromatin regions (Figure 6B).

When SMARCA5 is included in the model, the DMRT1–DNA interactions are maintained (Figure 6E), but the DNA near the DMRT1 motif is displaced farther from the histone core compared to the DMRT1-only model (Figure 6D, F), indicating chromatin loosening induced by SMARCA5. These findings suggest that while DMRT1 can recognize its target motifs in closed chromatin, it requires SMARCA5 to remodel nucleosomes and generate an accessible chromatin state.

### SMARCA5 establishes chromatin states that confer retinoic acid responsiveness

Finally, we sought to determine how *de novo* chromatin accessibility at distal DMRT1 binding sites contributes to spermatogenic differentiation. Loss of *Smarca5* impairs the transition from A_undiff_ to A_diff_, a process known to require retinoic acid (RA) signaling.^61,62^ Notably, the absence of A_diff_ in *Smarca5*-cKO testes phenocopies that observed in vitamin A (the RA precursor)–deficient mice, which lack RA.^62^ *Stra8*, a well-established RA-responsive gene, is first expressed in A_diff_ and later again in preleptotene spermatocytes.^63,64^ Retinoic acid receptors (RARs), members of the nuclear receptor family, act as ligand-dependent transcription factors.^65^ Supporting their role in RA signaling, pan-RAR ChIP-seq data from germline stem (GS) cells—an in vitro model of A_undiff_^66^—have shown RAR binding at the *Stra8* promoter.^40^

We therefore reanalyzed the pan-RAR ChIP-seq data^66^ and found that both SMARCA5 and DMRT1 also bind RAR-binding sites in A_undiff_ (Figure 7A). These RAR-binding regions were inaccessible in ProSG but became accessible in A_undiff_ in a SMARCA5-dependent manner (Figure 7B). For example, SMARCA5-dependent accessible chromatin was established at the promoters of *Stra8* and another RA-responsive gene, *Rec8*, both of which are co-occupied by DMRT1 and RAR (Figure 7C). These findings indicate that the SMARCA5–DMRT1 pioneering complex establishes a chromatin state that permits RAR binding, thereby priming A_undiff_ for RA-induced differentiation.

**Figure 7.**
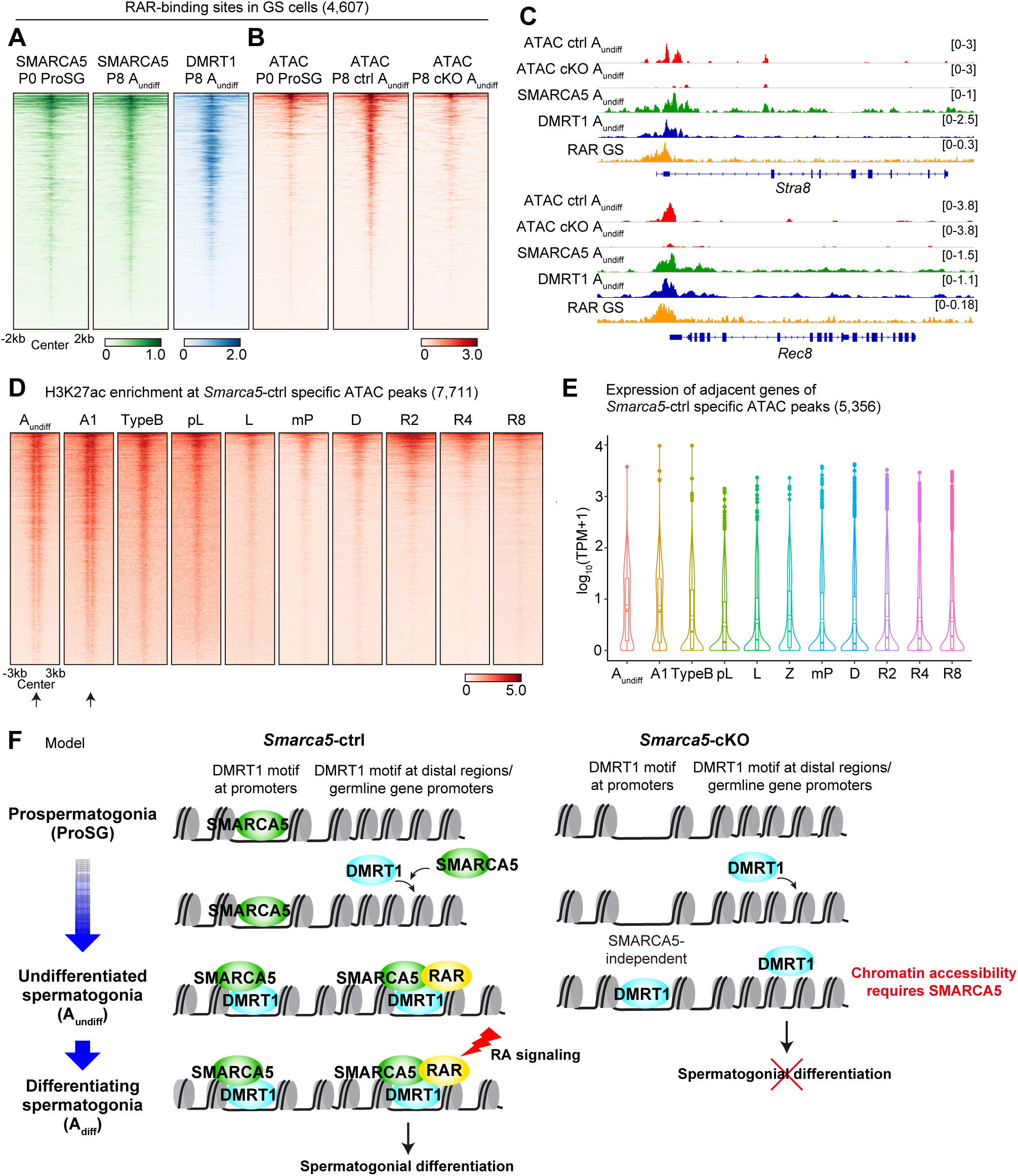
SMARCA5 shapes chromatin states that enable retinoic acid responsiveness. (A and B) Heatmaps showing SMARCA5 and DMRT1 enrichment (A), and ATAC enrichment (B) at RAR binding sites identified in GS cells (n = 4,607). (C) Track views of the *Stra8* and *Rec8* loci showing ATAC-seq signals, CUT&Tag profiles for SMARCA5 and DMRT1, and RAR ChIP-seq signals. (D) Heatmap showing H3K27ac enrichment at *Smarca5*-ctrl-specific ATAC peaks (n=7,701) during spermatogenesis, from A_undiff_ to round spermatids. Developmental stages: A1, Type A1 spermatogonia; TypeB, Type B spermatogonia; pL, preleptotene; L, leptotene; mP, mid-pachytene; D, diplotene; R2, steps 1–2 round spermatids; R4, steps 3–4; R8, steps 7–8. Arrows below the A_undiff_ and A1 panels indicate the absence of H3K27ac peaks at the center of ATAC-seq peaks. (E) Violin plots of RNA-seq expression (log_10_ (TPM+1)) for genes associated with *Smarca5*-ctrl-specific ATAC peak regions (n = 5,356). Z: zygotene spermatocytes. (F) Model summarizing SMARCA5 functions during the transition from ProSG to A_undiff_. SMARCA5 cooperates with DMRT1 to prime gene regulatory elements during the transition from ProSG to A_undiff_ and is essential for establishing a permissive chromatin environment for subsequent RAR recruitment. By doing so, SMARCA5 licenses transcriptional responses to retinoic acid (RA) that enable spermatogenic differentiation.

We reanalyzed the H3K27ac ChIP-seq data during spermatogenesis^14^ and found that SMARCA5-dependent accessible sites in A_undiff_ were enriched for H3K27ac in both A_undiff_ and A1 spermatogonia (a subset of A_diff_), and H3K27ac signals decreased upon further differentiation (Figure 7D). Notably, in A_undiff_ and A_diff_, H3K27ac signals were enriched in regions several hundred base pairs upstream and downstream of the ATAC-seq peak centers, while the peak centers themselves lacked H3K27ac (indicated by arrows in Figure 7D). This pattern likely reflects transcription factor binding at the central regions, flanked by H3K27ac-modified nucleosomes. Moreover, 5,356 genes located near these regions were highly expressed in A_undiff_ and A_diff_ (Figure 7E). These results support the notion that SMARCA5-dependent accessible sites act as enhancers in spermatogonia. In summary, SMARCA5 promotes spermatogonial differentiation by establishing chromatin accessibility at putative poised enhancers and germline gene promoters—including *Dmrt1*, *Stra8*, and *Rec8*—thereby priming the genome for RA responsiveness during the transition from ProSG to A_undiff_ (Figure 7F).

## Discussion

In this study, we report that SMARCA5 is an essential chromatin remodeler that cooperates with the pioneer transcription factor DMRT1 to prime gene regulatory elements during the transition from ProSG to A_undiff_ (Figure 7F). We demonstrate that, in SMARCA5-deficient spermatogonia, many enhancers and key germline gene promoters—including those of *Stra8* and *Rec8*—fail to acquire chromatin accessibility. Although DMRT1 is bound to many of these loci, the chromatin remains inaccessible, indicating that SMARCA5 is required to establish a permissive chromatin environment for subsequent recruitment of transcriptional regulators such as RARs. The failure to differentiate into A_diff_ in the absence of SMARCA5 suggests that disrupted epigenetic priming impedes RAR function. Thus, SMARCA5 confers developmental competence to male germ cells by establishing the chromatin states necessary for RA-responsive gene expression (Figure 7F).

Our finding that DMRT1, while capable of binding closed chromatin, requires SMARCA5 to generate full chromatin accessibility at many of its target sites (Figure 6) is a central conceptual advance. This observation refines the classical model of pioneer transcription factors, suggesting that in certain developmental contexts, their chromatin-opening activity depends on cooperation with chromatin remodelers. While in vitro nucleosome reconstitution assays have demonstrated that pioneer factors can open compact chromatin independently^28,67,68^, studies in cultured cells indicate that chromatin remodelers are often required to achieve chromatin accessibility^69,70^. Transcription requires the recruitment of various factors, including transcription factors and RNA polymerase II, a process that may involve the activity of chromatin remodelers to establish fully open chromatin regions^70^. Our findings provide *in vivo* evidence—using cells derived from living organisms rather than cultured cells—that chromatin remodeling is an essential step in the functional activation of pioneer factor-bound regions.

Additionally, DMRT6—expressed in A_diff_ and type B spermatogonia—binds genomic regions that largely overlap with those of DMRT1. These DMRT6 binding sites also became accessible in a SMARCA5-dependent manner during the transition from ProSG to A_undiff_ (Figure S5H), suggesting that DMRT6 functions sequentially after DMRT1 at shared regulatory sites.

Of note, a recent study also reported that *Smarca5*-cKO using a different *Ddx4*-Cre driver^71^ and the same *Smarca5*-floxed allele^35^ as in our study leads to male infertility.^72^ However, in that model, deletion of *Smarca5* did not lead to an acute and uniform loss of spermatogonia but allowed continued spermatogonial differentiation and meiotic progression.^72^ This discrepancy might be due to differences in Cre-induced recombination efficiency between the two *Ddx4*-Cre drivers, likely reflecting distinct levels of Cre expression. The *Ddx4*-Cre driver used in our study^36^ is a transgenic line with over 20 copies of the transgene inserted into the genome^73^, whereas the *Ddx4*-Cre driver^71^ used in the other study^72^ is a knock-in line. Incomplete loss of SMARC5 during early development in that model might have allowed spermatogonia to progress, thereby revealing that SMARCA5 is also required during later stages of spermatogenesis.^72^ Interestingly, SMARCA5 protein is present in the XY body of pachytene spermatocytes, a hallmark of meiotic sex chromosome inactivation (MSCI), an essential step in male meiosis^74^ (Figure S1C). Notably, our previous studies identified SMARCA5 in immunoprecipitation and mass spectrometry analyses of γH2AX-containing nucleosomes,^13,75^ which are enriched in the XY body. These findings suggest that SMARCA5 also functions during later stages of spermatogenesis beyond the A_diff_ stage.

In addition to its role in differentiation, we observe that SMARCA5 is also involved in maintaining the undifferentiated state in A_undiff_. This function may be mediated by PRC2-dependent H3K27me3, which is enriched at many distal SMARCA5 binding sites in these cells (Figure 4D). H3K27me3 is a hallmark of SSCs and is typically lost during the onset of differentiation, accompanied by activation of genes such as *Stra8.*^44^ During the transition from ProSG to A_undiff_, H3K27me3 is deposited at promoters already marked by H3K4me3, forming bivalent domains.^76^ At key loci such as *Dmrt1* and *Stra8*, these bivalent domains coincide with SMARCA5-dependent accessible regions, suggesting that SMARCA5 might facilitate the establishment of a poised chromatin state necessary for SSC maintenance. Thus, SMARCA5 may serve dual roles: one in Polycomb-mediated repression to maintain SSC identity and another in Polycomb-associated epigenetic priming that prepares cells for differentiation. These functions may not be mutually exclusive and likely occur in coordination with DMRT1. Accordingly, our study also clarifies the molecular function of DMRT1 in germ cells, highlighting a parallel to its role in somatic supporting cells (Sertoli cells) in the testes, where DMRT1 partners with Polycomb complexes to repress the female transcriptional program.^77^

Previous studies have shown that epigenetic states undergo substantial changes during the ProSG to A_undiff_ transition.^78,79^ Single-cell ATAC-seq analyses of developing testicular cells from E18.5 to P5.5 have revealed that newly accessible regions are enriched for binding motifs of various transcription factors, including DMRT1 and FOXO1^78^, a factor essential for spermatogonial maintenance.^80^ Notably, FOXO1 has been identified as a pioneer factor in cancer cells^81^. In light of our new findings, these observations suggest that additional regulators may contribute to establishing accessible chromatin during this developmental transition.

The current study further raises the intriguing possibility that the establishment of male epigenetic priming may suppress the female program during development from the bi-potential state of primordial germ cells, similar to the role of DMRT1 in Sertoli cells in the testes^82^. Notably, *Smarca5*-cKO A_undiff_-specific ATAC peak regions, which in the presence of SMARC5 remain inaccessible, were enriched for binding motifs shared by the transcription factors TCF3 and TCF12 (Figure 3E). TCF3 and TCF12 are primarily required for the female germline, particularly in supporting oocyte growth.^83^ Thus, it is tempting to speculate that loss of SMARCA5 may lead to the emergence of ectopic, female-like chromatin states.

Although pioneer transcription factors have been extensively studied in the context of early embryogenesis^84,85^ and cellular reprogramming,^86^ their roles during later developmental transitions remain poorly understood. Our findings highlight that the interplay between chromatin remodelers and pioneer factors is critical not only during embryogenesis but also during germline differentiation, where they enable transcriptional programs responsive to external signals. This finding expands the conceptual framework of epigenetic priming and underscores the broader importance of chromatin remodeling in shaping cellular competence across diverse developmental systems.

## Acknowledgements

We thank members of the Namekawa lab, Kanako Ikami, and Shosei Yoshida for stimulating discussions; Arthur I. Skoultchi for helping share *Smarca5^F/F^* mice; Azim Surani for providing *Stella*-GFP transgenic mice; So Maezawa for the homemade Tn5 transposase for ATAC-seq; Kei-Ichiro Ishiguro for the MEIOSIN antibody; and David Zarkower for the DMRT1 antibody. We acknowledge the following funding sources: JSPS Overseas Challenge Program for Young Researchers, TOYOBO Biotechnology Foundation, and JSPS Overseas Research Fellowship to Y.K.; and NIH Grant GM141085 to S.H.N.

## Author contribution

Y.K. and S.H.N. designed the study. Y.M. performed the RNA-seq experiments and ATAC-seq experiments. Y.K., Y.M., and M.H. performed the CUT&Tag experiments. Y.K., H.A., S.M., M.R., and S.P.K. performed staining experiments. H.A. performed Western blotting experiments. Y.K. performed the computational analysis, with the contribution from Y.M. Y.K. and S.O. performed structure prediction using AlphaFold3. Y.K., Y.M., H.A., M.H., S.O., D.J.P., R.M.S., and S.H.N. interpreted the results. D.J.P. provided the *Smarca5^F/F^* mouse line. Y.K., R.M.S., and S.H.N. wrote the manuscript with critical feedback from all authors. S.H.N. supervised the project.

## Methods

### Animals

Mice were maintained on a 12:12 light cycle in a temperature and humidity-controlled vivarium (22±2°C: 40-50% humidity) with free access to food and water in a pathogen-free animal care facility. Mice were used according to the guidelines of the Institutional Animal Care and Use Committee (protocol no. IACUC2018-0040, 21943, and 23545) at Cincinnati Children’s Hospital Medical Center and the University of California, Davis. *Smarca5-*cKO mice (*Smarca*5*^F/-^*; *Ddx4*-Cre [FVB-Tg (*Ddx4*-cre)1Dcas/J]) were generated from *Smarca5^F/F^* females crossed with *Smarca*5*^F/+^*; *Ddx4*-Cre males. *Smarca5*-ctrl in experiments were *Smarca5^F/+^*; *Ddx4*-Cre for the next generation sequencing experiments, and *Smarca5^F/+^*; *Ddx4*-Cre or *Smarca5^F/+^* for immunostaining and testis size measurements from littermates of *Smarca5* cKO. *Smarca5^F/F^* and *Ddx4*-Cre mouse lines were maintained on a background of FVB. *Smarca5^F/F^* mice were obtained from Dr. Davis J. Picketts,^35^ and *Ddx4*-Cre transgenic mice were purchased from the Jackson laboratory.^36^ *Stella*-GFP transgenic mice were obtained from Dr M. Azim Surani,^87^ maintained on a mixed genetic background of FVB and C57BL/6J.

### Histology and immunostaining

For the preparation of paraffin blocks, testes were fixed with 4% paraformaldehyde containing 0.05% Triton X-100 for 2 days at room temperature. Testes were dehydrated by a series of ethanol and then replaced with xylene and embedded in paraffin. For HE staining, 5 μm-thick paraffin sections were deparaffinized and stained with hematoxylin (Sigma, MHS16) and eosin (Sigma, 318906). For immunostaining, 5 μm-thick paraffin sections were deparaffinized and autoclaved in target retrieval solution (DAKO) for 10 min at 121°C. Sections were blocked with Blocking One Histo (Nacalai, 06349-64) for 20 min at room temperature and then incubated with primary antibodies overnight at 4°C. Sections were washed with PBST (PBS containing 0.1% Tween 20) three times at room temperature for 5 min and then incubated with the corresponding secondary antibodies. Finally, sections were counterstained with DAPI and mounted using 30 μL undiluted ProLong Gold Antifade Mountant (ThermoFisher Scientific, P36930). Images were obtained with an ECLIPSE Ti-2 microscope (Nikon) or BZ-X810 (Keyence).

### Western blotting

Testis pieces obtained from P7 testis were homogenized in RIPA buffer (50 mM Tris–HCl, pH 7.5; 150 mM NaCl; 0.1% SDS; 1% Triton X-100; 1% sodium deoxycholate) containing a protease inhibitor cocktail (Roche, 11697498001) and a phosphatase inhibitor cocktail (Sigma, P0044). For SMARCA5 detection, 20 μg of protein were separated by electrophoresis with a 10% SDS-PAGE gel, and the proteins were transferred using Trans-Blot® Turbo Transfer System (BIO-RAD) onto a PVDF membrane (EMD Millipore; IPVH00010). The membranes were blocked with StartingBlock™ T20 (TBS) Blocking Buffer (ThermoFisher Scientific, 37543) for 30 min at room temperature and then incubated with primary antibodies overnight at 4°C. After being washed with TBST three times, membranes were then incubated with secondary antibodies conjugated to HRP (Abcam, ab131366 or ab131368) for 1 h at room temperature, and bands were visualized using an ECL kit according to the manufacturer’s instructions (EMD Millipore; WBKLS0500).

### Flow cytometry and cell sorting

Flow cytometric experiments and cell sorting were performed using SH800S (SONY), with antibody-stained testicular single-cell suspensions prepared as described previously^44,88^ with minor modifications. Briefly, to prepare single cell suspensions for cell sorting, detangled seminiferous tubules were incubated in 1× Krebs–Ringer Bicarbonate Buffer (Sigma, K4002) supplemented with 1.5 mg/ml collagenase Type 1 and 0.04 mg/ml DNase I at 37°C for 15 min with gentle agitation and dissociated using vigorous pipetting, and then add 0.75 mg/ml hyaluronidase (Sigma, H3506) and incubated at 37°C for 10 min with gentle agitation and dissociated using vigorous pipetting. The cell suspension was centrifuged at 300 × g, the cells resuspended in 10 ml FACS buffer (PBS containing 2% FBS), and then centrifuged again at 300 × g for 5 min, and this step was repeated one more time. The pelleted cells were resuspended in 1 ml FACS buffer and then filtered through a 70 μm nylon cell strainer (Falcon, 352350). The resultant single cells were stained with cocktails of antibodies diluted with FACS buffer, listed as follows: PE-conjugated anti-mouse/human CD324 (E-cadherin) antibody (1:500, Biolegend, 147303), PE/Cy7-conjugated anti-mouse CD117 (c-Kit) antibody (1:200, Biolegend, 105814), and FITC-conjugated anti-mouse CD9 antibody (1:500, Biolegend, 124808). After a 50-minute incubation on ice, cells were washed with 10 ml FACS buffer three times by centrifugation at 300 × g for 5 min and filtered into a 1 ml FACS tube through a 35 μm nylon mesh cap (Falcon, 352235). 7-AAD Viability Stain (Invitrogen, 00-6993-50) was added to the cell suspension for the exclusion of dead cells. Samples were kept on ice until sorting. Cells were analyzed after removing small and large debris in FSC-A versus SSC-A gating, doublets in FSC-W versus FSC-H gating, and 7AAD^+^ dead cells. Then the desired cell population was collected in gates determined based on antibody staining.

### RNA-seq library generation and sequencing

RNA-seq libraries of A_undiff_ from *Smarca5*-ctrl and cKO and A_diff_ from *Smarca5*-ctrl were prepared as described;^88^ briefly, 10,000 A_undiff_ or A_diff_ cells were pooled from two independent mice as one replicate, and two independent biological replicates were used for RNA-seq library generation. Total RNA was extracted using the RNeasy Plus Micro Kit (QIAGEN, Cat # 74034) according to the manufacturer’s instructions. Library preparation was performed with NEBNext® Single Cell/Low Input RNA Library Prep Kit for Illumina® (NEB, E6420S) according to the manufacturer’s instructions. Prepared RNA-seq libraries were sequenced on the HiSeq X Ten system (Illumina) with paired-ended 150-bp reads.

### ATAC-seq library generation and sequencing

ATAC-seq libraries of germ cells were prepared as described;^88^ briefly, 10,000 A_undiff_, A_diff_, or P0 prospermatogonia (ProSG) were pooled from two independent mice as one replicate. Samples were lysed in 50 μl of lysis buffer (10 mM Tris–HCl (pH 7.4), 10 mM NaCl, 3 mM MgCl2, and 0.1% NP-40, 0.1% Tween-20, and 0.01% Digitonin) on ice for 5 min. Immediately after lysis, the samples were spun at 500 × g for 10 min at 4°C, and the supernatant was removed. The sedimented nuclei were then incubated in 10 μl of transposition mix (0.5 μl homemade Tn5 transposase (∼1 μg/μl), 5 μl 2× tagment DNA buffer (10 mM Tris–HCl (pH 7.6), 10 mM MgCl2, and 20% dimethyl formamide), 3.3 μl PBS, 0.1 μl 1% digitonin, 0.1 μl 10% Tween-20, and 1 μl water) at 37°C for 30 min in a thermomixer with shaking at 500 rpm. After tagmentation, the transposed DNA was purified with a MinElute kit (Qiagen). Polymerase chain reaction (PCR) was performed to amplify the library using the following conditions: 72°C for 3 min; 98°C for 30 s; thermocycling at 98°C for 10 s, 60°C for 30 s, and 72°C for 1 min. Quantitative PCR was used to estimate the number of additional cycles needed to generate products at 25% saturation. Seven to eight additional PCR cycles were added to the initial set of five cycles. Amplified DNA was purified by 1.0x SPRIselect beads (Beckman Coulter). ATAC-seq libraries were sequenced on the HiSeq X Ten system with 150-bp paired-end reads.

### CUT&Tag library generation and sequencing

CUT&Tag libraries of ctrl A_undiff_ and ProSG for SMARCA5 were prepared as previously described (a step-by-step protocol https://www.protocols.io/view/bench-top-cut-amp-tag-kqdg34qdpl25/v3) using CUTANA™ pAG-Tn5 (Epicypher, 15-1017)^89^. Quantitative spike-in CUT&Tag for DMRT1 of *Smarca5*-ctrl and cKO A_undiff_ was performed by adding *Drosophila* S2 cells at a 1:5 ratio to mouse spermatogonial cells (5,000 S2 cells to 25,000 mouse cells) in each reaction. The antibodies used were rabbit anti-SMARCA5 (1/100) or rabbit anti-DMRT1 antibody (1/50). CUT&Tag libraries were sequenced on the Novaseq X Plus system with 150-bp paired-end reads.

### RNA-seq data processing

Raw paired-end RNA-seq reads after trimming by Trim-galore (https://github.com/FelixKrueger/TrimGalore) (version 0.6.7) were aligned to the mouse (GRCm38/mm10) genome using STAR^90^ (version STAR_2.5.4b) with following options: --outSAMtype BAM SortedByCoordinate; --twopassMode Basic; --outFilterType BySJout; --outFilterMultimapNmax 1; --winAnchorMultimapNmax 50; --alignSJoverhangMin 8; --alignSJDBoverhangMin 1; --outFilterMismatchNmax 999; --outFilterMismatchNoverReadLmax 0.04; --alignIntronMin 20; -- alignIntronMax 1000000; --alignMatesGapMax 1000000 for unique alignments. To quantify aligned reads in RNA-seq, aligned read counts for each gene were generated using featureCounts^91^ (v2.0.1), which is part of the Subread package based on annotated genes (gencode.vM25.annotation.gtf)^92^. The transcripts per million (TPM) values of each gene were used for comparative expression analyses and computing the Pearson correlation coefficient between biological replicates using corrplot.

To detect differentially-expressed genes (DEGs) between *Smarca5*-ctrl A_undiff_ and *Smarca5*-ctrl A_diff_, or *Smarca5*-ctrl A_undiff_ and *Smarca5*-cKO A_undiff_, DESeq2^93^ (version 1.42.1) was used for differential gene expression analyses with cutoffs ≥2-fold change and binomial tests (Padj < 0.05; P-values were adjusted for multiple testing using the Benjamini–Hochberg method). Padj values were used to determine significantly dysregulated genes.

GO term analysis was performed using the website tool DAVID (https://david.ncifcrf.gov/home.jsp)^94,95^. GO term was visualized by ggplot2 (version 3.4.4) of the R package based on gene number, fold enrichment, and *P* value. A violin plot was drawn using the R package ggplot2.

### ATAC-seq and CUT&Tag data processing

Raw paired-end ATAC-seq reads after trimming by Trim-galore were aligned to either the mouse (GRCm38/mm10) genomes using bowtie2 (version 2.3.3.1)^96^ with default arguments. Raw paired-end CUT&Tag reads after trimming by Trim-galore were aligned to either the mouse (GRCm38/mm10) genomes using Bowtie2 (version 2.3.3.1) with options: –end-to-end –very-sensitive –no-mixed –no-discordant –phred33 -I 10 -X 700. All unmapped and non-uniquely mapped reads were filtered out by samtools (version 1.9)^97^ before being subjected to downstream analyses. PCR duplicates were removed using the ‘MarkDuplicates’ command in Picard tools (version 2.23.8) (https://broadinstitute.github.io/picard/, Broad Institute). For CUT&Tag on DMRT1, *D. melanogaster* DNA delivered by Drosophila S2 cells was used as spike-in DNA, as described.^89^ For mapping *D. melanogaster* spike-in fragments, we also aligned to either the *D. melanogaster* (dm6) genome using Bowtie2 and used the ‘–no-overlap –no-dovetail’ options to avoid cross-mapping using Bowtie2. PCR duplicates were removed using the ‘MarkDuplicates’ command in Picard tools. Spike-in normalization was implemented using the exogenous scaling factor computed from the dm6 mapping files (scale factors = 10000/spike-in reads for DMRT1 CUT&Tag).

Biological replicates were pooled for visualization and other analyses after validation of reproducibility. Peak calling for ATAC-seq data was performed using MACS3 (version 3.0.0a7)^98^ with the parameters: -g mm --nomodel --nolambda. Peak calling for CUT&Tag data was performed using MACS2 (version 2.2.7.1) with the parameters: -g mm. We computed the number of overlapping peaks between peak files using BEDtools (version 2.28.0) ^99^ function intersect. To detect genes adjacent to ATAC-seq and CUT&Tag peaks, we used the HOMER (version 4.9.1)^100^ function annotatePeaks.pl. The deeptools was used to draw tag density plots and heatmaps for read enrichment. To visualize ATAC-seq and CUT&Tag data on SMARCA5 using the Integrative Genomics Viewer (Broad Institute),^101^ bins per million (BPM) normalized counts data were created from sorted BAM files using the deeptools. To visualize CUT&Tag data on DMRT1, spike-in normalized genome coverage tracks with 1 bp resolution in BigWig format were generated using ‘bamCoverage’ from deepTools (version 3.5.5)^102^ with the parameters ‘--binSize 1 --extendReads --samFlagInclude 64 --normalizeUsing RPKM –scaleFactor $scale_factor’.

### AlphaFold3 modeling

AlphaFold 3^57^ was run using the AlphaFold web server (https://alphafoldserver.com). Two amino acid sequences translated from *Hist1h2ao, Hist1h2bb*, *H3f3a*, and *Hist2h4* were used as inputs to form a histone core. The sequence of the 601 sequence with 60 bp of flanking DNA and DMRT1 motif sequence is as follows:

5’−CTGGAGAATCCCGGTGCCGAGGCCGCTCAATTGGTCGTAGACAGCTCTAGCACCGCTTAAACGCACGTACGCGCTGTCCCCCGCGTTTTAACCGCCAAGGGGATTACTCCCTAGTCTCCAGGCACGTGTCAGATATATACATCTTGATACATTGTATCACAGCGACCTTGCCGGTGCCAGTCGGATAGTGTTCCGAGCTCCCACTCT−3’

ChimeraX (version 1.9) was used for visualization.

### ChIP-seq data reanalysis

Raw single-end H3K27ac ChIP-seq data were downloaded from Gene Expression Omnibus (GEO) under accession no. GSE132446 and GSE242515. Fastq files of biological replicates were merged and then trimmed by Trim-galore. Trimmed reads were aligned to the mouse (mm10) and Drosophila (dm6), respectively, using bowtie2 (version 2.3.3.1) with the parameters: --very-sensitive --phred33. PCR duplicates were removed using the ‘MarkDuplicates’ command in Picard tools (version 2.23.8). Spike-in normalization was implemented using the exogenous scaling factor computed from the dm6 mapping files (scale factors = 1000000/spike-in reads). Deeptools of the bamCoverage program was used to generate spike-in normalized coverage tracks (bigwig format) with the parameters: --binSize 10 -- normalizationUsing RPGC --extendReads --effectiveGenomeSize 2652783500 –scaleFactor $scaleFactor.

### Statistics

Statistical methods and *P* values for each plot are listed in the figure legends and/or in the Methods. Next-generation sequencing data (RNA-seq, CUT&Tag) were based on two independent replicates. No statistical methods were used to predetermine sample size in these experiments. Experiments were not randomized, and investigators were not blinded to allocation during experiments and outcome assessments.

## Data availability

ChIP-seq data for H3K27ac in A_undiff_ (THY1^+^) spermatogonia were downloaded from GSE130652. ChIP-seq data for H3K4me3 in A_undiff_ spermatogonia were downloaded from GSE89502. CUT&Tag data for H3K27me3 and H2AK119ub in A_undiff_ spermatogonia were downloaded from the GSE221944. ChIP-seq data for H3K27ac in male germline were downloaded from GSE132446 and GSE242515. RNA-seq data in the male germline were downloaded from GSE242515. ChIP-seq data for RAR in GS cells were downloaded from GSE1116798. ChIP-seq data for DMRT6 in adult testis were downloaded from GSE60440.

## Code availability

Source code for all software and tools used in this study, with documentation, examples, and additional information, is available at the URLs listed above.

