## Supplementary material for "The SMARCA5–DMRT1 Pioneer Complex Establishes Epigenetic Priming to Direct Male Germline Development": Figures S1-S5

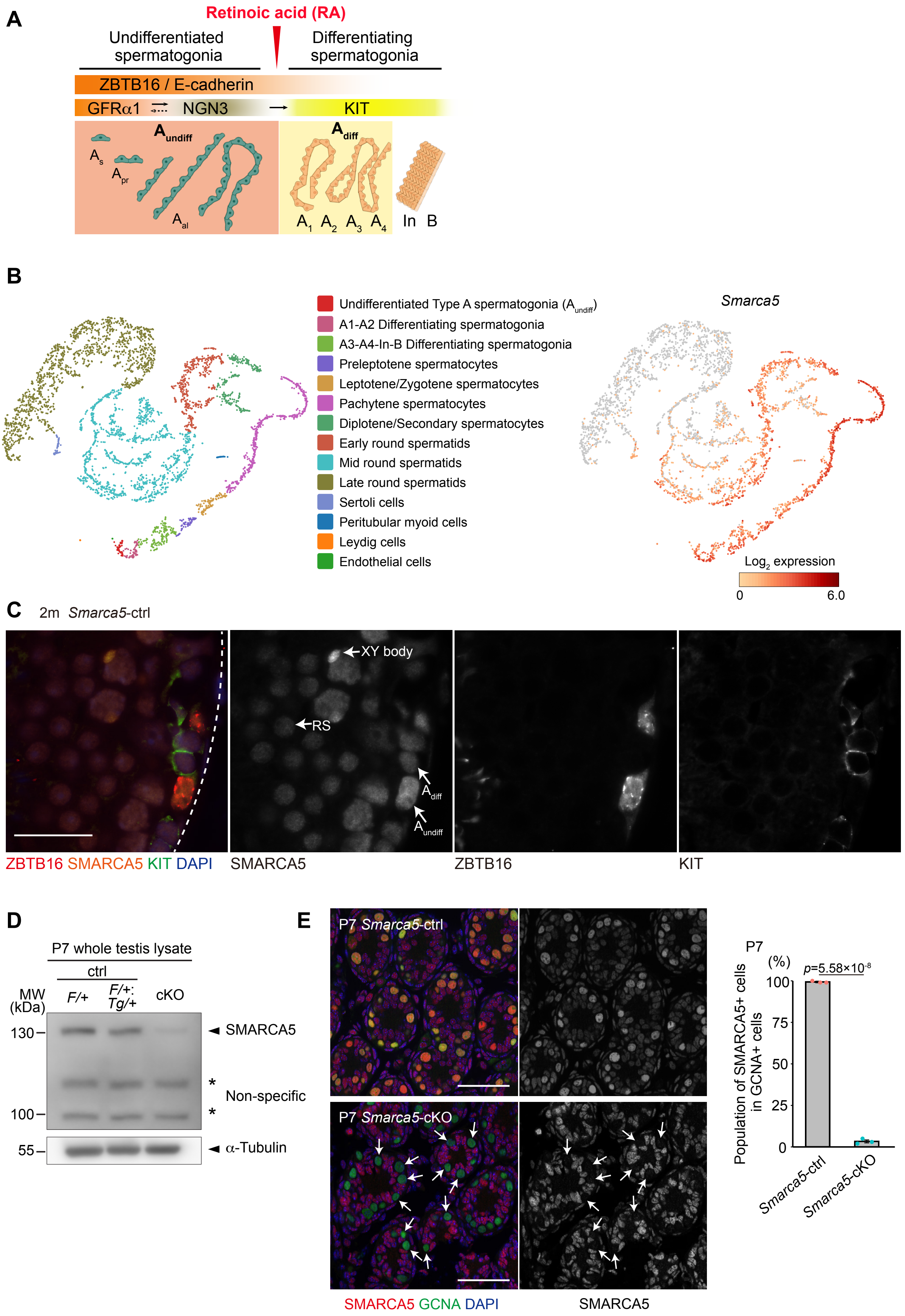

A
Retinoic acid (RA)
Undifferentiated spermatogonia
Differentiating spermatogonia
ZBTB16 / E-cadherin
GFRα1
NGN3
KIT
A undiff
A diff
A s
A pr
A al
A 1 A 2 A 3 A 4
In B
B
Undifferentiated Type A spermatogonia (A undiff)
A1-A2 Differentiating spermatogonia
A3-A4-In-B Differentiating spermatogonia
Preleptotene spermatocytes
Leptotene/Zygotene spermatocytes
Pachytene spermatocytes
Diplotene/Secondary spermatocytes
Early round spermatids
Mid round spermatids
Late round spermatids
Sertoli cells
Peritubular myoid cells
Leydig cells
Endothelial cells
Smarca5
Log2 expression
0
6.0
C
2m Smarca5-ctrl
XY body
RS
A diff
A undiff
ZBTB16 SMARCA5 KIT DAPI
SMARCA5
ZBTB16
KIT
D
P7 whole testis lysate
ctrl
MW (kDa)
F/+
F/+; Tg/+
cKO
130
100
55
SMARCA5
Non-specific
α-Tubulin
E
P7 Smarca5-ctrl
P7 Smarca5-cKO
SMARCA5 GCNA DAPI
SMARCA5
P7
(%)
p=5.58×10-8
100
75
50
25
0
Population of SMARCA5+ cells in GCNA+ cells
Smarca5-ctrl
Smarca5-cKO

**Figure S1. SMARCA5 expression during spermatogenesis and its deletion in the germline**
(A) Schematic representation of the spermatogonial hierarchy and marker gene expression (created with BioRender.com). Spermatogonia are classified as type A, intermediate (In), or type B. Type A includes undifferentiated spermatogonia ($A_{undiff}$, including $A_{single}$ ($A_s$), $A_{paired}$($A_{pr}$) and $A_{aligned}$($A_{al}$)), and early differentiating spermatogonia ($A_{diff}$, including $A_1$, $A_2$, $A_3$ and $A_4$), while In and type B represent later stages of differentiating spermatogonia.
(B) t-SNE plot of single-cell RNA-seq data from adult testis,[35] showing identification of cell clusters (left) and *Smarca5* expression across clusters (right).
(C) Immunostaining of SMARCA5, ZBTB16, and KIT in 2-month-old *Smarca5*-ctrl testes. In the SMARCA5-stained panel, representative $A_{undiff}$, $A_{diff}$, XY body, and round spermatid (RS) are indicated by arrows. Scale bar, 25 µm.
(D) Western blot analysis of testicular lysate obtained from *Smarca5*$^{F/+}$, *Smarca5*$^{F/+}$; *Ddx4*-Cre$^{+}$ (*Smarca5*-ctrl) and *Smarca5*$^{F/F}$; *Ddx4*-Cre$^{+}$ (*Smarca5*-cKO) at P7. Asterisks indicate the positions of non-specific bands.
(E) Testis sections of Smarca5-cKO and control littermates at P7 stained with DAPI and antibodies against SMARCA5 and GCNA (germ cell marker). Scale bar, 100 µm. The graph shows the percentage of SMARCA5$^{+}$ cell in GCNA$^{+}$ cells. Error bars represent mean ± s.d. 100 GCNA$^{+}$ cells for each samples were counted. Independent values obtained from individual mice are shown as dots. SMARCA5$^{-}$GCNA$^{+}$ cells are shown with arrows. Statistical significance was assessed using a two-tailed unpaired Student's t-test assuming equal variances (n = 3 per group).

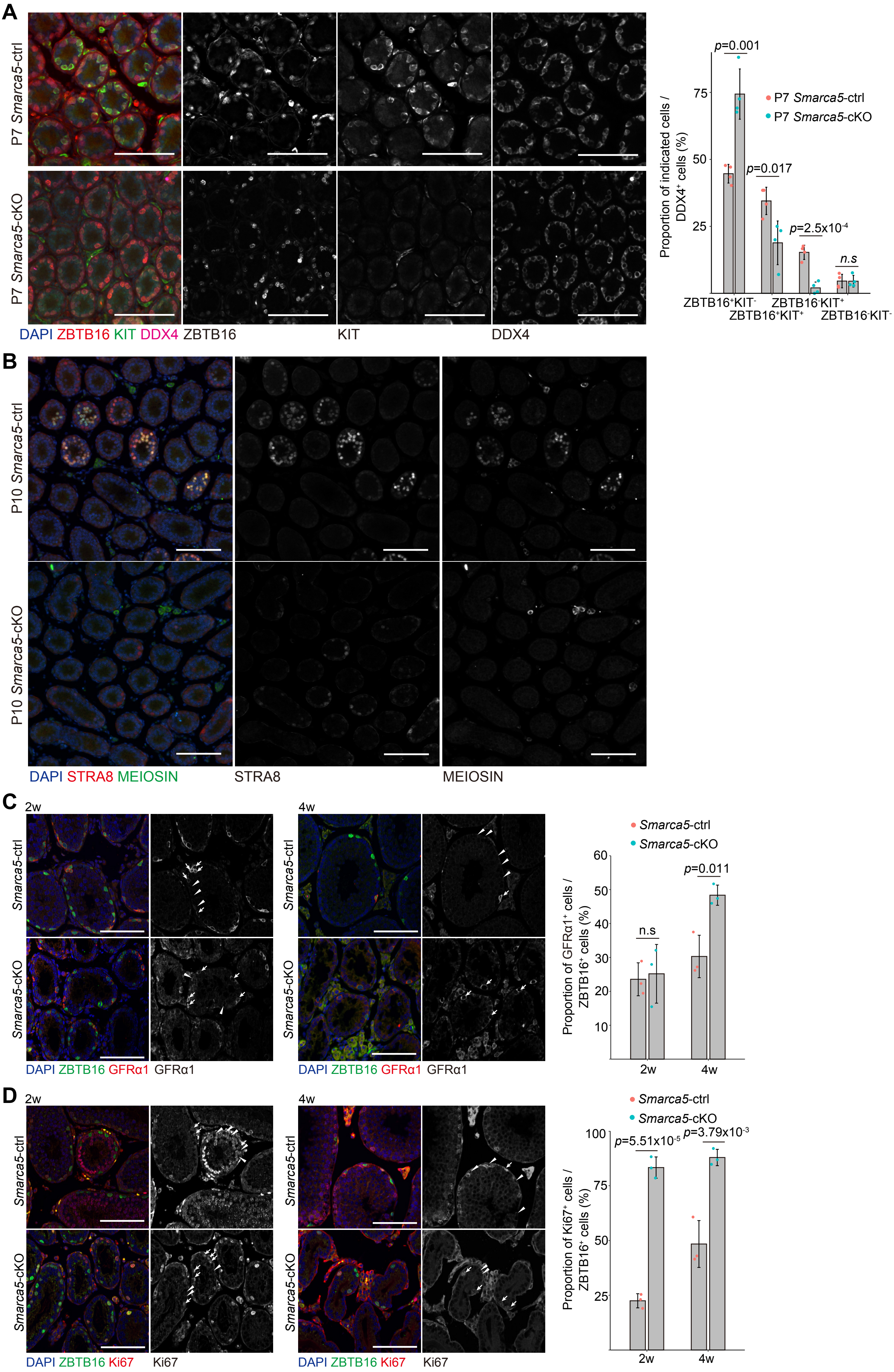
A
P7 Smarca5-ctrl
P7 Smarca5-cKO
DAPI ZBTB16 KIT DDX4
ZBTB16
KIT
DDX4
Proportion of indicated cells / DDX4+ cells (%)
p=0.001
p=0.017
p=2.5x10-4
n.s
P7 Smarca5-ctrl
P7 Smarca5-cKO
ZBTB16+KIT-
ZBTB16+KIT+
ZBTB16-KIT+
ZBTB16-KIT-
B
P10 Smarca5-ctrl
P10 Smarca5-cKO
DAPI STRA8 MEIOSIN
STRA8
MEIOSIN
C
2w
4w
Smarca5-ctrl
Smarca5-cKO
DAPI ZBTB16 GFRα1
GFRα1
Proportion of GFRα1+ cells / ZBTB16+ cells (%)
n.s
p=0.011
D
DAPI ZBTB16 Ki67
Ki67
Proportion of Ki67+ cells / ZBTB16+ cells (%)
p=5.51x10-5
p=3.79x10-3

**Figure S2. Immunohistochemical analysis of *Smarca5*-cKO testis sections.**

(A) Testis sections from *Smarca5*-cKO and control littermates at P7 stained with DAPI and antibodies against ZBTB16, KIT, and DDX4. Scale bars, 100 µm. The graph shows the percentage of ZBTB16$^+$ and/or KIT$^+$ cells among DDX4$^+$ germ cells. Error bars represent mean ± s.d. At least 180 DDX4$^+$ germ cells were counted per sample. Data points represent values from individual mice. Statistical significance was assessed using a two-tailed unpaired Student's t-test assuming equal variances (n = 4 per group).

(B) Testis sections from *Smarca5*-cKO and control littermates at P10 stained with DAPI and antibodies against STRA8 and MEIOSIN (a marker of preleptotene spermatocytes). Scale bars, 100 µm.

(C and D) Testis sections from *Smarca5*-cKO and control littermates at 2 and 4 weeks of age stained with DAPI and antibodies against ZBTB16 and Ki67 (C) or GFRα1 (D). Scale bars, 100 µm. Graphs show the percentage of Ki67$^+$ (C) or GFRα1$^+$ (D) cells among ZBTB16$^+$ cells. Error bars represent mean ± s.d. At least 120 ZBTB16$^+$ cells were counted per sample. Data points represent individual mice. Statistical significance was determined using a two-tailed unpaired Student's t-test assuming equal variances (n = 3 per group). Ki67$^+$ and GFRα1$^+$ cells are indicated by arrows; negative cells are indicated by arrowheads.

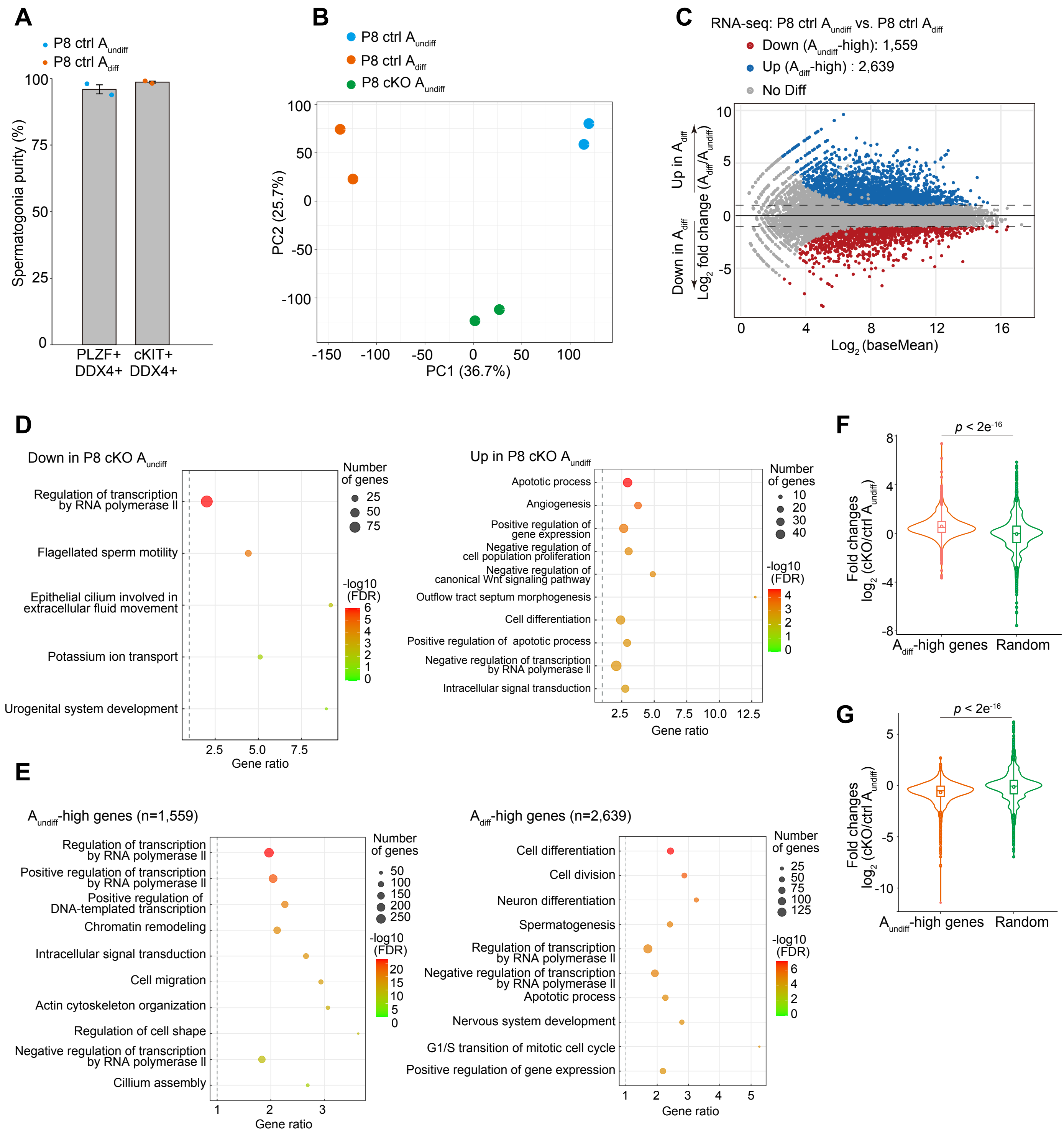


**Figure S3. RNA-seq analysis of $A_{undiff}$ and $A_{diff}$.**

(A) Quantification of the purity of isolated spermatogonia. Purity values from two independent experiments are shown as individual dots.

(B) Principal component analysis (PCA) plot showing biological replicates of RNA-seq samples.

(C) Transcriptomic comparison between $A_{undiff}$ and $A_{diff}$ populations. Differentially expressed genes (DEGs) are defined as those with $Log_2$ fold change > 2, *Padj* < 0.05, based on a binomial test with Benjamini-Hochberg correction.

(D) Gene Ontology (GO) analysis of genes downregulated (left) and upregulated (right) in *Smarca5*-cKO $A_{undiff}$ compared to *Smarca5*-ctrl $A_{undiff}$.

(E) GO analysis of genes downregulated ($A_{undiff}$-high genes: left) and upregulated ($A_{diff}$-high genes: right) in *Smarca5*-ctrl $A_{diff}$ compared to ctrl $A_{undiff}$.

(F and G) Violin plots showing $log_2$ fold change in *Smarca5*-cKO $A_{undiff}$ compared to ctrl $A_{undiff}$ within gene groups highly expressed in $A_{diff}$ ($A_{diff}$-high genes: n = 2,639; F) or $A_{undiff}$ ($A_{undiff}$-high genes: n = 1,559; G). For comparison, $log_2$ fold change values from randomly selected genes are also shown. Box plots indicate the 25th, median, and 75th percentiles; dots within the box represent the mean. Statistical significance was based on Levene's test and Wilcoxon rank sum test.

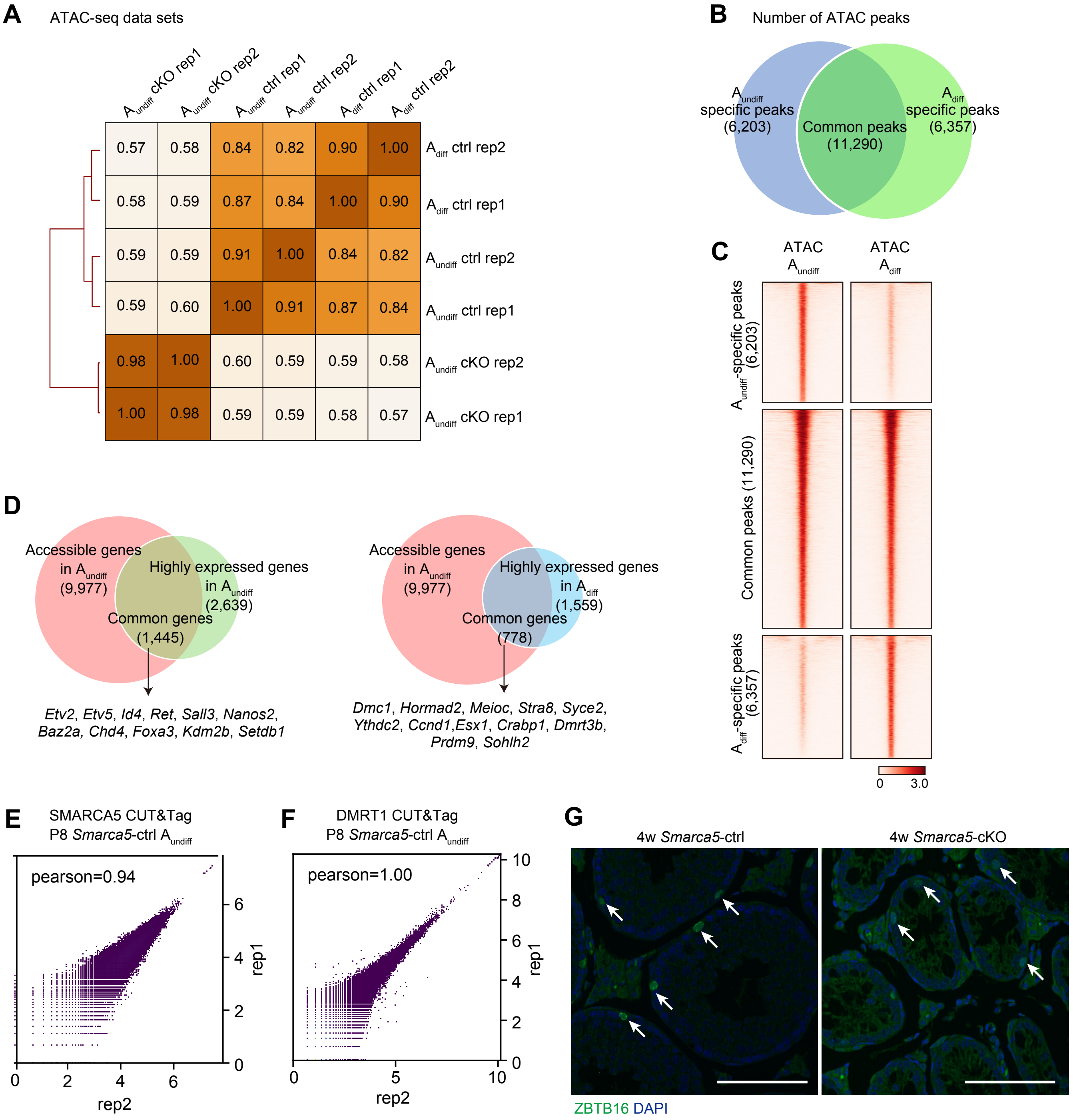

**Figure S4. ATAC-seq and CUT&Tag analysis of $A_{undiff}$ and $A_{diff}$.**
(A) Heatmap showing Pearson correlation coefficients among each cell type based on ATAC-seq profiles. Two biological replicates were merged for downstream analysis.
(B) Venn diagram showing the overlap of ATAC-seq peaks between $A_{undiff}$ and $A_{diff}$.
(C) Heatmaps showing ATAC-seq enrichment in $A_{undiff}$ and $A_{diff}$ at $A_{undiff}$-specific, common, or $A_{diff}$-specific peaks.
(D) Venn diagrams showing the overlap between accessible genes in $A_{undiff}$ (n = 9,977) and genes highly expressed in $A_{undiff}$ compared to $A_{undiff}$ (left), or highly expressed in $A_{diff}$ compared to $A_{diff}$ (right). Genes located within ±10 kb of ATAC-seq peaks in $A_{undiff}$ were defined as accessible.
(E and F) Scatter plots showing Pearson correlation between two biological replicates for SMARCA5 CUT&Tag (E) and DMRT1 CUT&Tag (F) in P8 *Smarca5*-ctrl $A_{undiff}$. Two replicates were merged for downstream analysis.
(G) Testis sections from *Smarca5*-cKO and control littermates at 4 weeks stained with DAPI and antibodies against ZBTB16. Arrows indicate ZBTB16$^{+}$ cells. Scale bars, 100 µm.

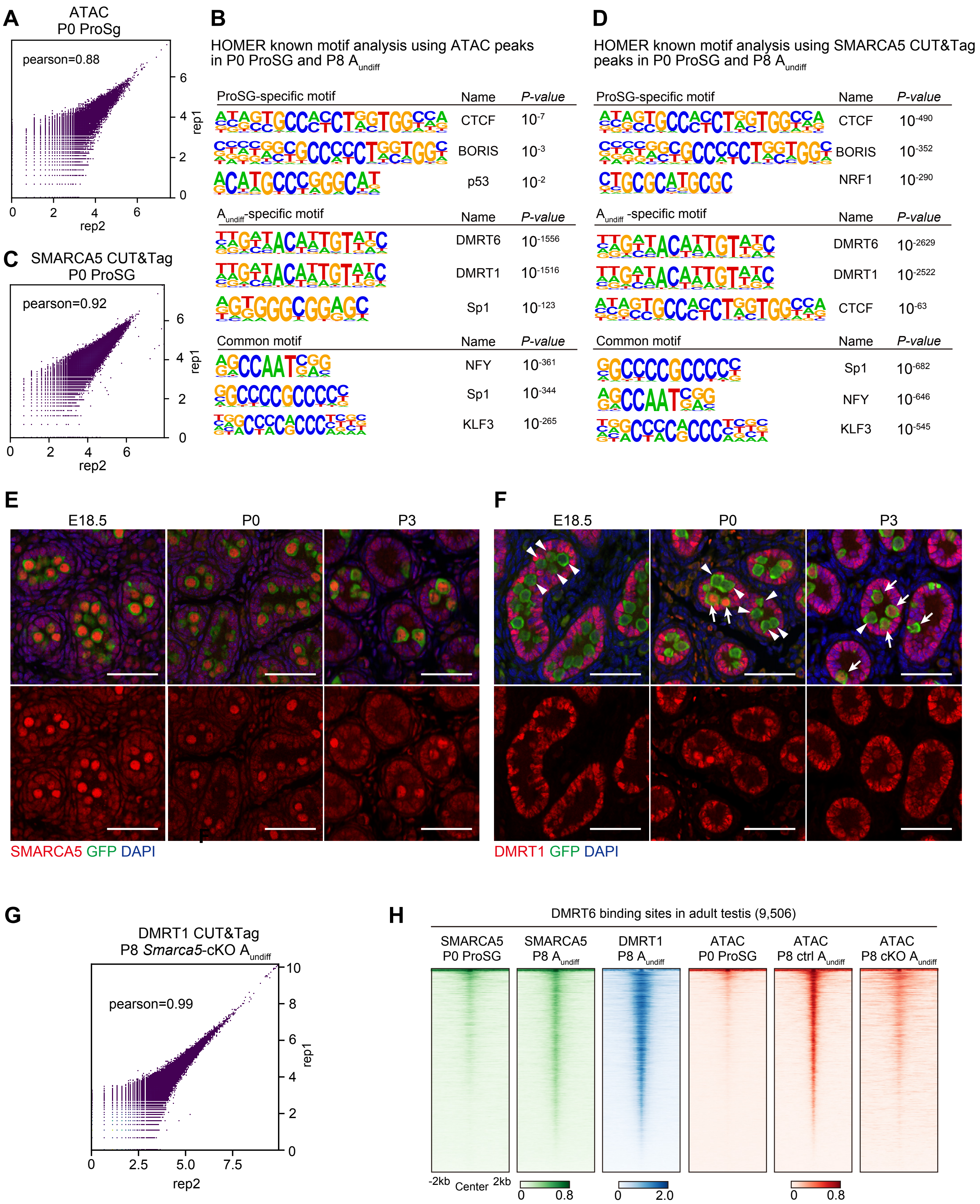
A
ATAC
P0 ProSg
pearson=0.88
rep1
rep2
B
HOMER known motif analysis using ATAC peaks in P0 ProSG and P8 A$_{undiff}$
ProSG-specific motif | Name | P-value
CTCF $10^{-7}$
BORIS $10^{-3}$
p53 $10^{-2}$
A$_{undiff}$-specific motif | Name | P-value
DMRT6 $10^{-1556}$
DMRT1 $10^{-1516}$
Sp1 $10^{-123}$
Common motif | Name | P-value
NFY $10^{-361}$
Sp1 $10^{-344}$
KLF3 $10^{-265}$
C
SMARCA5 CUT&Tag
P0 ProSG
pearson=0.92
rep1
rep2
D
HOMER known motif analysis using SMARCA5 CUT&Tag peaks in P0 ProSG and P8 A$_{undiff}$
ProSG-specific motif | Name | P-value
CTCF $10^{-490}$
BORIS $10^{-352}$
NRF1 $10^{-290}$
A$_{undiff}$ -specific motif | Name | P-value
DMRT6 $10^{-2629}$
DMRT1 $10^{-2522}$
CTCF $10^{-63}$
Common motif | Name | P-value
Sp1 $10^{-682}$
NFY $10^{-646}$
KLF3 $10^{-545}$
E
E18.5
P0
P3
SMARCA5 GFP DAPI
F
E18.5
P0
P3
DMRT1 GFP DAPI
G
DMRT1 CUT&Tag
P8 *Smarca5*-cKO A$_{undiff}$
pearson=0.99
rep1
rep2
H
DMRT6 binding sites in adult testis (9,506)
SMARCA5 P0 ProSG
SMARCA5 P8 A$_{undiff}$
DMRT1 P8 A$_{undiff}$
ATAC P0 ProSG
ATAC P8 ctrl A$_{undiff}$
ATAC P8 cKO A$_{undiff}$
-2kb
Center
2kb

**Figure S5. ATAC-seq and CUT&Tag analysis of ProSG and $A_{undiff}$.**
(A and C) Scatter plot showing Pearson correlation between two biological replicates of ATAC-seq (A) and SMARCA5 CUT&Tag (C) in P0 ProSG. Replicates were merged for downstream analysis.
(B and D) HOMER known motif analysis of ATAC-seq peaks (B) and SMARCA5 CUT&Tag peaks (D) at ProSG-specific, $A_{undiff}$-specific, and common sites.
(E and F) Testis sections from *Stalla*-GFP mice stained with DAPI and antibodies against GFP and SMARCA5 (E) or DMRT1 (F) at E18.5, P0, and P3. $DMRT1^{+}$ germ cells are indicated by arrows; $DMRT1^{-}$ cells are indicated by arrowheads. Scale bars, 100 μm.
(G) Scatter plot showing Pearson correlation between two biological replicates of DMRT1 CUT&Tag in P8 *Smarca5*-cKO $A_{undiff}$.
(H) Heatmap showing SMARCA5, DMRT1, and ATAC enrichment in ProSG and $A_{undiff}$ at DMRT6-binding sites identified in the adult testis (n = 9,506).
